# Multidecadal changes in land cover across a disturbance gradient in mountain grasslands of Kyrgyzstan

**DOI:** 10.64898/2026.03.24.712710

**Authors:** Sherry C.E. Young, Hannah V. Watkins, Steven Brownlee, Helen F. Yan, Isabelle M. Côté

## Abstract

Mountain ecosystems face unprecedented pressures from anthropogenic activities and climate change, challenging the productivity of these vital habitats. In the Tien Shan mountains, understanding localized responses to these pressures is often hindered by the coarse spatiotemporal resolutions of available data. To address this, we combined high-resolution satellite imagery (1997-2021) to map land-cover dynamics in the Naryn oblast, Kyrgyzstan across a gradient of grazing intensities. We classified and quantified land-cover distribution over 24 years, investigating the roles of topography, elevation, and anthropogenic disturbances as drivers of change. Our results identify intermediate elevations, high degrees of disturbance, and the interaction between the two as the primary contributors to recent transitions in grassland, forest, and barren habitats. By integrating Landsat analysis-ready data, European Space Agency WorldCover dataset and digital elevation models at fine spatial scales, we provide valuable contemporary and historical landscape and habitat-level insights and a high-resolution framework for disentangling climate-driven shifts from land-use impacts. These findings highlight the urgency of localized management in remote, data-poor regions where rapid environmental change threatens both biodiversity and pastoral livelihoods. Our work serves as a critical baseline for characterizing the adaptability of semi-arid mountain rangelands under escalating global and regional pressures.

## 1. Introduction

Approximately one-third of the world’s forests and terrestrial biodiversity, including half of the world’s biodiversity hotspots, are on mountains (Chape et al., 2008; Gratzer & Keeton, n.d.; Grumbine & Xu, 2021). Mountains cover up to one-quarter of the Earth’s landmass beyond Antarctica. While the definition of a mountain varies depending on local perceptions, conceptions, and culture (Smith & Mark, 2003) the most accepted defining features are ruggedness, vertical atmospheric pressure and temperature gradients (Körner, 2007; Körner et al., 2021). The interaction between such heterogeneous, complex terrain and climate creates a high diversity of land cover, microclimate and therefore habitat types. Mountain habitats also boast high levels of endemic species (Körner, 2021; Noroozi et al., 2018). Considered the world’s water towers (Immerzeel et al., 2020; Viviroli et al., 2007), mountains provide fresh water for over half of the global human population (Egan & Price, 2017), as well as myriad opportunities for resource production and extraction, recreation, education, spirituality, and cultural heritage (Bernbaum, 2022; Canedoli et al., 2024). At a smaller scale, mountain habitats support local economies based on farming and herding. Mountain communities, especially in the highlands, are isolated and have few resources, and thus are particularly vulnerable to changes in climate (i.e., extreme events, shifting seasons) and environmental conditions (i.e., habitat degradation, pollution). With unprecedented rates of global human expansion and climate change over the last several decades, people worldwide have observed the rapid pace at which alpine regions are changing.

Mountain ecosystems continue to experience particularly rapid physical and environmental change due to anthropogenic global warming (Gerrard, 1991; Marston, 2008). Warmer temperatures, shifts in precipitation regimes and increased frequency of extreme events seriously challenge the persistence of contemporary mountain habitats and both human and ecological communities (Keiler et al., 2024; Knight, 2022). The reported impacts of climate change in mountain areas include increased natural hazards (Terzi et al., 2019) and decreasing snow cover, permafrost area and glacier mass balance (Immerzeel et al., 2020). For instance, glaciers retreated in the 2010s at almost twice the pace as they did in the 1980s (Pelto et al., 2020). Retreating and melting glaciers jeopardize water availability for people and ecosystems existing in already arid environments. In addition, while decreases in snow cover and glaciers expose more land with the potential to provide new suitable habitat, these darker areas have lower albedo and thus absorb solar radiation, leading to warmer land surfaces (Zhang et al., 2022). Climate change also drives shifts in ecosystem composition and plant migration (Kelly & Goulden, 2008; Parmesan & Yohe, 2003). Cold-adapted alpine plant communities are highly sensitive to warming (Inouye, 2020), and temperature increases across elevation gradients result in the loss of endemic and cold-adapted species, species movement from low to high elevations, increased competition and overall changes in community composition and ecosystem functioning (Freeman et al., 2018). Finally, climate-change-induced impacts are amplified by the history, intensity and scale of land use practices, leading to accelerated land degradation in a growing number of cases (Orr et al., 2024). While impacts of warming and land use on high-altitude ecosystems are often discussed in general terms at a global scale, it remains a challenge to predict how local and/or regional mountain areas will respond due to the structural heterogeneity and complexity of these vast, remote, and rugged landscapes as well as the political and governance history of the region (Reyer et al., 2017).

The Tian Shan Mountain range, which extends from China to Uzbekistan, provides an example of drastic recent changes in land use, linked to political upheaval. Called the *celestial* or *heavenly mountain*, this extensive system of primarily arid and semi-arid mountain ranges covers approximately 1,000,000 km^2^, including peaks of more than 7,000 m and low-lying areas >150 m below sea level. As part of the Palaearctic steppes (Wesche et al., 2016), the Tian Shan Mountains’ arid and semi-arid grasslands provide one of the richest source of livelihood for humans. In Kyrgyzstan, the most mountainous country in Middle Asia (> 90% of land area is mountain; FAO, 2015), rangelands constitute most agricultural lands, making Kyrgyzstan a great example of how mountain habitats can economically support local communities through farming and herding. Historically, nomadic herders moved across the landscape, from low elevations in winter to highlands in summer, known as vertical transhumance. This vertical transhumance enabled access to forage for livestock throughout the seasons without the exhaustion of natural resources (Aryal et al., 2018). These ancient patterns were severely disrupted in the 1920s following annexation by Russia and incorporation into the Soviet Union, and then again following the collapse of the USSR in 1991, which saw an abrupt decline in livestock numbers (Chi et al., 2020).

However, over the last 25 years, reduced transhumance and the increasing numbers of livestock across rangelands have led to a growing concern about rangeland productivity. While observations of land degradation are not unanimous throughout the country (Levine et al., 2019), an expanding body of evidence from herders and scientists is raising concerns about pasture degradation (Eddy et al., 2017; Hoppe et al., 2016; Zhumanova et al., 2018a). Land degradation is defined as a temporary or permanent decline in a system’s productive capacity and potential to be used, including for economic purposes (Stocking, 2001). The increasing amount of livestock and the loss of movement through transhumance leads to concentrated grazing pressures resulting in pasture degradation, also referred to as browning (Cook & Pau, 2013), which differs from natural seasonal senescence. In 2022, pasture degradation maps showed that 81.5% of winter pastures were severely degraded (vs < 43% in other seasons; Domenech et al., 2022). This alarming degradation rate is forcing herders to supplement feed their herds, jeopardizing the sustainability of local communities already relying on animal herding as a subsidence-based livelihood.

High rates of pasture degradation also pose a major concern for ecosystem functioning. Plant growth depends on nitrogen mineralization and soil organic carbons that increase with rising precipitation (Burke & Rundquist, 2021; Liu, Zhou, et al., 2021; Wilsey, 2018). Arid and semi-arid grasslands have limited, periodic water availability, extreme temperature seasonality and short growing seasons, which has selected for adaptations such as small growth forms, high root-shoot ratio, reduced leaf area, and a more efficient photosynthetic pathway (C3) (Taft et al., 2011). As a result, mountain grassland ecosystems typically have low net primary productivity (NPP) (Wingler & Hennessy, 2016), are particularly sensitive to environmental change, and recover slowly from disturbances. Significant warming rate in Central Asia over the last 30 years (+0.4 °C; Ma et al., 2021), extreme climatic events and ongoing pressures from land use (i.e., grazing activities) have drastically altered grassland dynamics, structure and functioning (Yang et al., 2023; Zhao et al., 2023), leading to gradual degradation and potentially permanent state conversion (Zhang et al., 2018; Zhang et al., 2018). Changes in grassland functioning and structure have cascading effects on other trophic levels and species interactions, such as soil dynamics and microorganisms and herbivory through reduced foraging, which can ultimately alter predator-prey dynamics that involve emblematic mammals, such as the snow leopard (*Panthera uncia*), argali (*Ovis ammon*) and Siberian ibex (*Capra sibirica)*. It is unclear if degraded pastures are now permanently altered, highlighting the urgency of assessing and restoring grasslands before it is too late.

Given the established importance of mountain ecosystems through the myriads of ecosystem services they provide and the increasing intensity of challenges they face, it is crucial to assess potential patterns of rangeland degradation at the fine scales at which management can be implemented. Using satellite imagery, we i) classified, quantified, and mapped land cover distribution between 1997 and 2021 at fine spatio-temporal resolutions and ii) compared land cover distribution between areas with different conservation statuses in the Naryn oblast, the region with the greatest concentration of grasslands in Kyrgyzstan (Figure 1). These adjacent conservation designations offer a gradient of anthropogenic pressures from areas with prohibited herding to areas with intense herding. Then, we investigated the role of (1) topography in driving the spatial distribution of land cover classes in the region and (2) anthropogenic disturbances and elevation (as a proxy for global warming) as drivers of changes in land cover over time. While we expected elevation to be the most important driver of land cover distribution (Han et al., 2024; Schrodt et al., 2019; Setiawan et al., 2024), we also predicted that the combination of slope and aspect might play a role (Borchardt et al., 2011; Yang et al., 2020) along with ruggedness as a measure of habitat heterogeneity (Jay et al., 2023; Malanson et al., 2023). The widespread use of sensor data with large spatial resolutions (e.g., 500 m or 1 km) often limits the characterization of land cover change within more localized and smaller areas. The fine-scale, high resolution methods used in this study provide a framework to disentangle the roles of topography, anthropogenic pressures and climate change in the distribution of land cover in an area with limited baseline assessment of plant diversity and assemblages.

**Figure 1.**
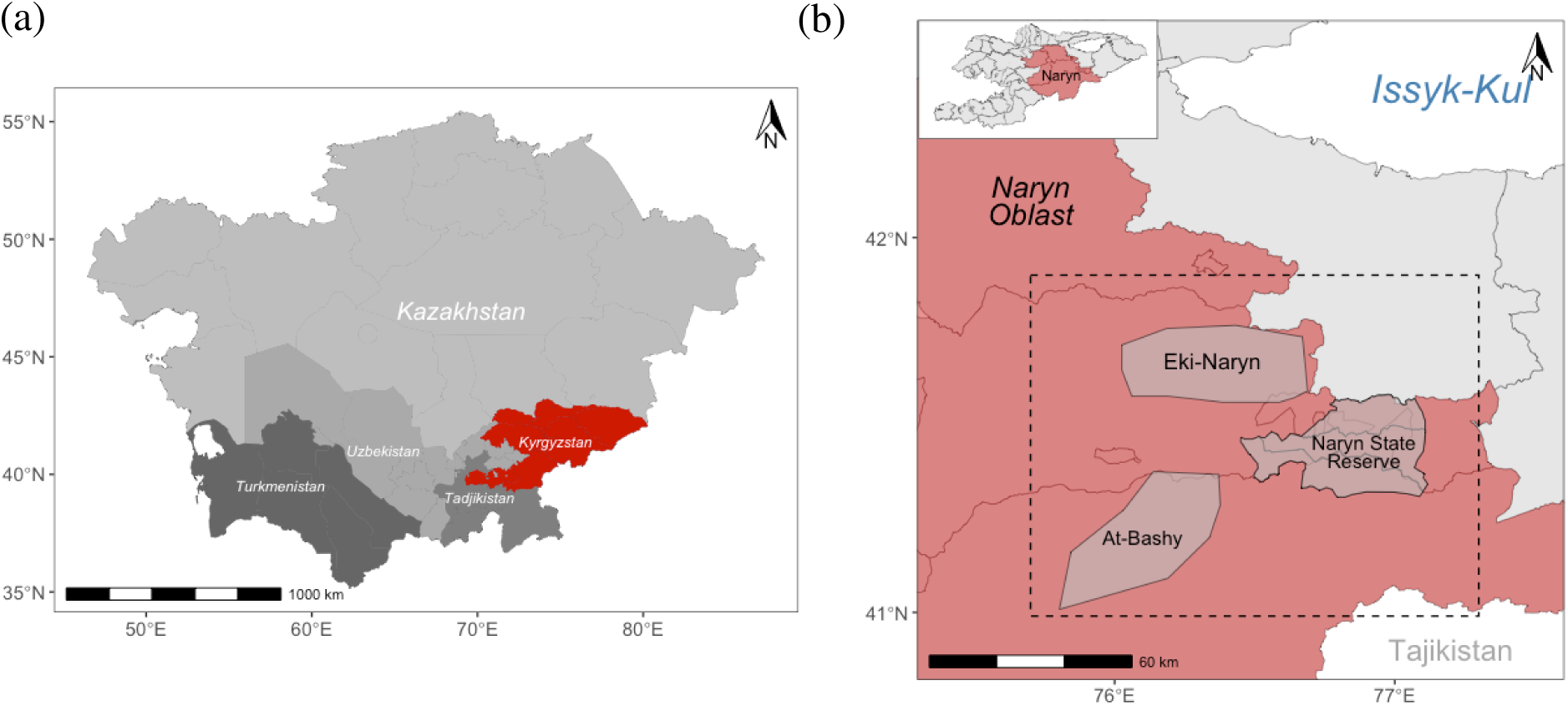
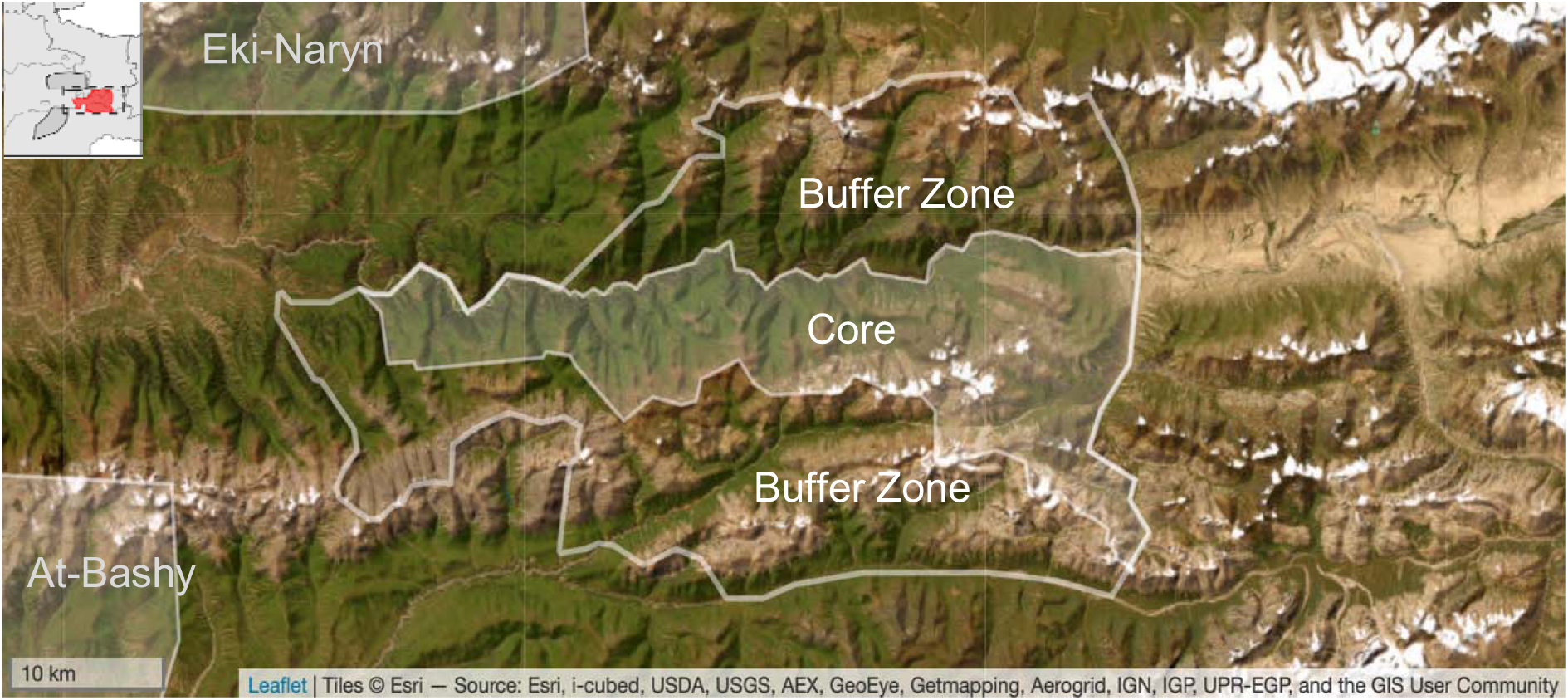
(a) Location of Kyrgyzstan (red) in Middle Asia. (b) The study areas in Kyrgyzstan’s Naryn oblast (red) include the Naryn State Reserve, Eki-Naryn Area and At-Bashy areas. Figure 0 Location of the core and buffer zones of the Naryn State Reserve.

## 2. Materials and Methods

### 2.1. Study area

The study took place in the Naryn oblast, the poorest of the seven administrative regions of Kyrgyzstan (Figure 1; Crewett, 2012). Located south-east of Kyrgyzstan, the Naryn oblast has the largest proportion of agricultural land in the country, with animal herding as the main source of livelihood (Azarov et al., 2020). Home to grassland valleys and tundra plateaux of the Inner Tian Shan, the region has a climate characterized by dry, cold, long winters and short warm and dry summers (Weather Spark, 2025), with daily mean temperatures ranging from -15.8 and 18.2 between January and August along with 290 mm of annual mean precipitation mainly occurring between April and July (Eddy et al., 2017). Naryn, the capital of the region, is considered the tenth largest city in Kyrgyzstan, with 41,000 inhabitants in 2021 (National Statistical Committee of the Kyrgyz Republic, 2023). We selected three study areas, each located in adjacent parts of the Inner Tian Shan range, that present similar topographic and landscape features but that together represent a gradient of anthropogenic pressures based on the history of grazing through local herding.

#### 2.1.1. Protected area and mild disturbances: Naryn State Reserve

The Naryn State Reserve (41°25′50″N, 76°35′11″E) is a 1,080 km^2^ reserve created in 1972 as a multi-use sanctuary (*zakaznik*), with a ban on hunting to protect elk populations (*Cervus elaphus*) (Farrington, 2005). In 1983, the sanctuary became a state nature reserve (*zapovednik*), which further enforced regulations to ensure the preservation, conservation and restoration of the ecosystem while providing research and educational opportunities (Farrington, 2005; Kozhokulov et al., 2021). The reserve presents two types of management: a fully protected core zone located south of the Naryn riverbanks and two large multi-use buffer zones located north and south of the core zone (Figure 2). The core zone ranges from 2379 m to 4470 m in altitude, and it is mostly characterized by north-facing slopes only accessible by rangers and research teams. Therefore, the core zone is considered as one end of the disturbance gradient, with minimal to no anthropogenic disturbances over the past 40 years. The buffer zones, however, have historically been shaped by intense grazing and forestry. Signs of degradation and soil erosion from this era were still visible in the early 2000s (Farrington, 2005), and the buffer zones continue to be used for subsidence-based herding to this day (OSI-Panthera, *personal communication*). The Tian Shan Mountain ash and Schrenk’s spruce constitute most of the forest cover, along with grasslands, barren lands and glaciers as main habitat types. Activities in the buffer zones are monitored, and the degree of disturbance is considered mild.

**Figure 2.**
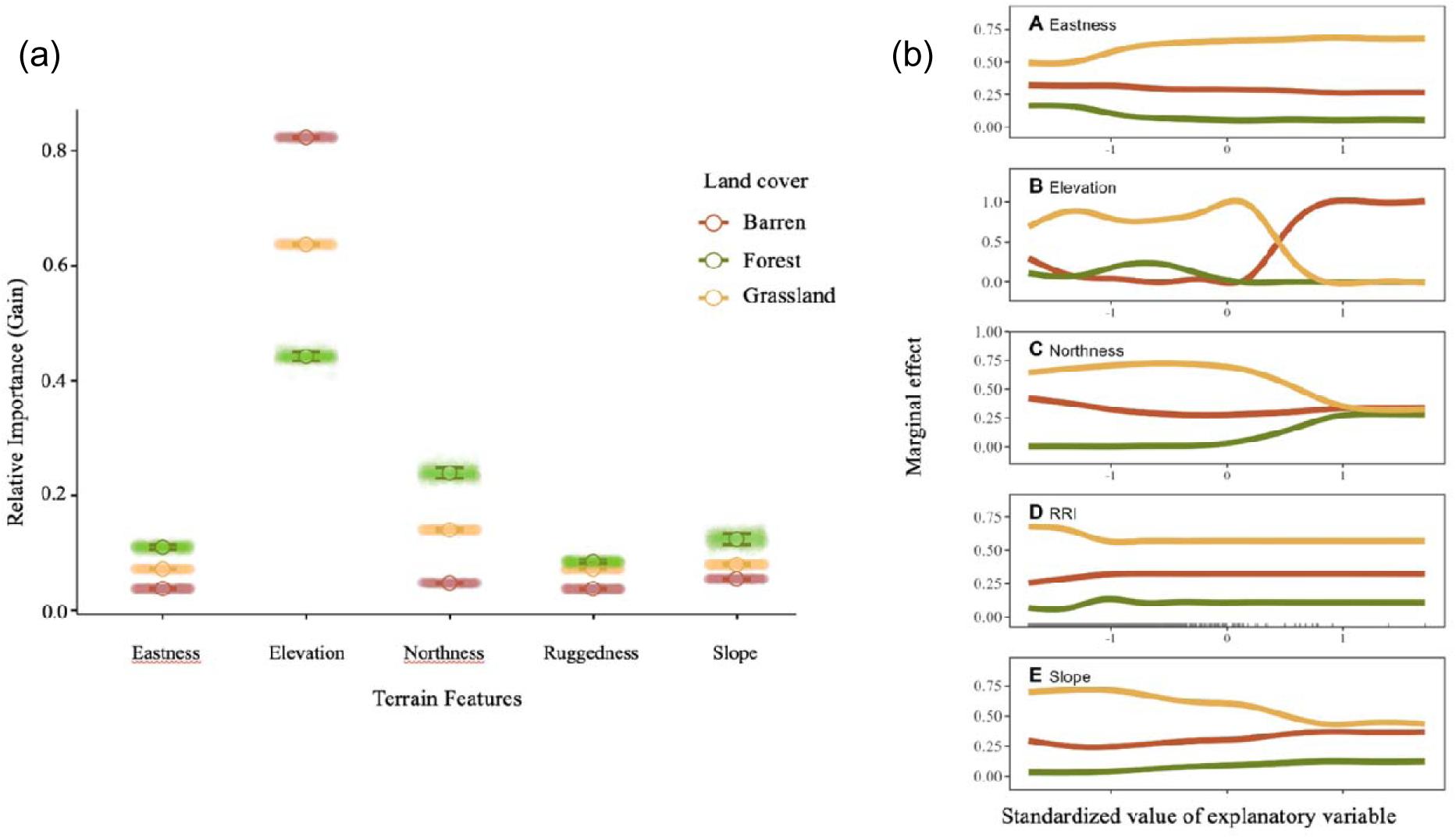
(a) Relative importance, expressed as gain, with a higher gain representing greater relative importance of the variable, of five terrain features in explaining the distribution of each land cover class after 1,000 boosted regression tree models. (b) Explanatory variables include A - eastness, B - elevation, C - northness, D - ruggedness (RRI) and E - slope (in degrees).

#### 2.1.2. Moderate disturbances: Eki-Naryn

The Eki-Naryn area is characterized by the small village of Eki-Naryn, which results in a moderate degree of disturbances compared to the Naryn State Reserve. Eki-Naryn is a small village located in an area with features like those of the Naryn State Reserve (i.e., surface area, elevation, slope, and aspect ranges), 8 km to the northwest (Figure 1b). The Eki-Naryn area ranges from 2314 m to 4422 m in elevation with the same characteristic continental, dry and seasonal climate. Overlapping with the Issyk-Kul region, humans from the time of nomads over 3,000 years used this area for herding and as a travel route. More recently, tourism has been growing in the region because of various exceptional landscape features such as the Eki Naryn valley (41°40’48.43”N, 76°25’27.40”E). Due to the proximity to human settlements and accessibility, this area is considered moderately disturbed.

#### 2.1.3. High disturbances: At-Bashy

The last study area, At-Bashy, represents a “highly disturbed” state. The At-Bashy Mountain range borders south of At-Bashy, one of the major cities in the Naryn oblast and capital of the At-Bashy district. This southern district of Naryn faces more drastic overgrazing than the Eki-Naryn area because of its closer proximity to villages. Recent maps indicate that pastures located in the At-Bashy range are already considered degraded (Domenech et al., 2022). In 2010, the At-Bashy district included 13,300 households (Azarov et al., 2020), from which 72 households were randomly sampled to understand the socio-economic situation of the local population. Each herder owned, on average, 3.85 ha of agricultural land and 15.8 livestock, with a strong preference for sheep, horse and cattle, and goats (Azarov et al., 2020). Despite an overall poor irrigation system, most of the land is used to produce fodder as supplementary feeding for livestock during the winter, which remains nutritionally insufficient. Sheep and horse herds migrate to higher pastures (*jailoo*) from June to September before returning near the settlement for the rest of the year, where cows stay throughout the year. Our defined study area ranges from 2072 m to 4396 m, south of the city of At-Bashy and is considered highly degraded due to the high level of disturbance (Domenech et al., 2022).

### 2.2. Geospatial data

#### 2.2.1. ESA land cover map

To classify and map the land cover in the Naryn region, we used the 2021 v200 European Space Agency (ESA) WorldCover 10 m resolution (Zanaga et al., 2022). The ESA WorldCover is a global land cover dataset that includes 11 land covers derived from the United Nations Land Cover Classification System (UN-LCCS): tree cover, shrubland, grassland, cropland, built-up, bare/sparse vegetation, snow and ice, permanent water bodies, herbaceous wetland, mangroves, moss and lichen. The ESA imagery uses both Sentinel-1 and Sentinel-2 satellite data. This combination of datasets allows mapping areas with frequent cloud cover, which is a common characteristic of high mountain areas. We used Google Earth Engine (Gorelick et al., 2017) to access the ESA imagery, after which we clipped the cloud-hosted tiles to our areas of interest and downloaded them. Then, we used the ESA land cover classification to perform a supervised classification of the Landsat composites (Appendix: Figure A1).

#### 2.2.2. Landsat data source

To characterize and track changes in land cover between 1997 and 2021, we used the open source 16-day 30 m Landsat Analysis Ready Data (ARD) provided by the Global Land Analysis and Discovery (GLAD) team at the University of Maryland (Potapov et al., 2020). The Landsat ARD provides normalized surface reflectance from 1997 to the present by combining and harmonizing Landsat 5 (™), 7 (ETM+) and 8 (OLI/TIRS) collection 2 (Table 1). We downloaded the Landsat ARD imagery from 1 January 1997 to 18 December 2021, using GLAD Tools, a user-friendly software package developed by the GLAD team to allow users to conduct various analyses and pre-processing steps to the ARD (Potapov et al., 2020). As three ARD tiles straddled our study areas, they were mosaiced together and clipped to the shape of each study area across the disturbance gradient. Land cover assessment and imagery for the 1990s are limited in Kyrgyzstan due to gaps in ground station coverage between the fall of the Soviet Union and the mid-2000s. As a result, imagery for some of this period is missing. Thus, we selected a target year and used the previous years (n = 4) annual Landsat imagery to fill in the gaps and create a multi-year composite at the expense of a finer temporal resolution. The aggregation was done for six time periods - 1997 to 2001, 2001 to 2005, 2005 to 2009, 2009 to 2013, 2013 to 2017, and 2017 to 2021, later referred to as 2001, 2005, 2009, 2013, 2017 and 2021 periods - to best classify and quantify land cover change. Each composite pixel represents the median value recorded in each pixel across a given period in each of four spectral bands - near-infrared (NIR), shortwave infrared 1 (SWIR1), shortwave infrared 2 (SWIR2) and red.

**Table 1.** Total predicted percentages of browning and greening across areas with various degrees of disturbance (area), the five 5-year time-periods (year), and elevation bands (elevation).The significance of the difference is reported through p-values. Results for each variable are marginalized across levels of the other variable (e.g., the proportion of cells undergoing browning in the core is averaged across years and elevations). To improve readability, we colored cells with percentages significantly higher than their counterpart. Cells with significantly higher browning are colored in red, and those with significantly higher greening are colored in yellow. Cells included percentages that are not significantly different are left blank.

| Variable | Level | Browning | Greening | p-value of pairwise comparison in transition type |
| --- | --- | --- | --- | --- |
| Area | Core | 3.5 % | 2.4% | 0.011 |
|  | Buffer | 2.7 % | 1.9% | 0.007 |
|  | Eki-Naryn | 4.1 % | 2.5% | 0.001 |
|  | At-Bashy | 2 % | 1.4% | 0.013 |
| Year | 2005 | 3.7 % | 2.3 % | 0.001 |
|  | 2009 | 4.2 % | 1.9 % | < 0.001 |
|  | 2013 | 2.3 % | 1.9 % | 0.155 |
|  | 2017 | 1.4 % | 2.4 % | 0.011 |
|  | 2021 | 3.5 % | 1.5 % | < 0.001 |
| Elevation | Band 1 | 3.6 % | 2 % | 0.003 |
|  | Band 2 | 2.7 % | 1.7 % | 0.021 |
|  | Band 3 | 3.8 % | 2.8 % | 0.084 |
|  | Band 4 | 2 % | 1.3 % | 0.013 |
|  | Band 5 | 0.7 % | 0.3 % | 0.031 |

Using a composite with shortwave bands and a composite with pseudo-natural colors, we compared which composite increases the accurate detection of vegetation cover types. Here, we used multi-year metric composites generated by GLAD Tools composed of NIR, SWIR1 and SWIR2 bands, which is expected to be more accurate in predicting vegetation types due to the presence of both shortwave infrared band (Li et al., 2022; Potapov et al., 2020) compared to our second composite composed of NIR, red and SWIR bands, more commonly used to predict land cover types (Potapov et al., 2020) (Appendix: Figure A1).

#### 2.2.3. Topographic variables

To quantify the topographic complexity of the region, we used the 30-m NASA Shuttle Radar Topography Mission Global (SRTMGL1 v003) digital elevation model (DEM) (NASA JPL, 2013). The SRTM data was downloaded using the GLAD Tools software (<u>GLAD TOOLS v2</u>; Potapov et al., 2020), thus providing an exact spatial overlap and match in resolution with the Landsat ARD. Slope, i.e., the degree of change in elevation in the direction of the water flow line, and aspect, i.e., the angular direction that a slope faces (Amatulli et al., 2020), are essential in creating topo climates in alpine landscapes. We generated slope and aspect, both in radians, as well as slope in degrees, by using the SRTM DEM rasters for each study area. Due to the aspect being a circular variable, we calculated eastness and northness to provide a continuous measure indicating a site’s north/south and west/east position (Amatulli et al., 2020). Northness and eastness are calculated as the sine of the slope in radians multiplied by the cosine and sine of aspect, respectively (Amatulli et al., 2020). To characterize the landscape’s ruggedness, we also included the radial roughness index (RRI), as a measure of terrain heterogeneity to predict species distribution across the landscape (Trevisani et al., 2023) (Appendix: Figure A1).

### 2.3. Data processing

#### 2.3.1. Plot selection

Each study area was separated into five elevation bands based on their respective elevation range, with a fixed width of 414 m (core zone), 428 m (buffer zones), 421 m (NW), and 465 m (SN) (Appendix: Figure A2). We reclassified the DEM for each elevation band to limit demands for computational power (Appendix: Table A1). We randomly extracted 1% of the pixels available within each band, with each pixel with a spatial resolution of 30 m (0.00025° x 0.00025°), producing a total sample size of 14,647 pixels for the buffer zones (439,410 m^2^) and 7997 pixels for the core zone (239,910 m^2^) of the Naryn State Reserve, 23,118 pixels for the Eki-Naryn area (693,540 m2) and 23,146 pixels for the At Bashy area (694,380m2) (Appendix: Figure A3, Table A2).

#### 2.3.2. Land Cover Classification

##### a. Reference data

We used the 2021 ESA WorldCover to characterize the land cover for each randomly generated sample site. First, we aggregated the 10m resolution of the ESA composite into a 30m resolution composite, defining the most frequent land cover class out of the 3 aggregated pixels as the main land cover. In the event of ties (i.e., 3 different classes), the pixel was removed.

Then, we classified each generated sample site, which were used as the training dataset to classify the Landsat multitemporal metric composites for each period at a spatial resolution of ∼ 30 m (0.00025° x 0.00025°). According to the 2021 ESA world cover dataset, more than half of our study area is characterized by grasslands, followed by 20% of barren land with sparse vegetation and moss and lichens (Appendix: Figure A4). Grasslands, bare/sparse vegetation and moss and lichens have similar surface reflectance signatures, leading to separability and characterization issues during the supervised classification. Thus, we decided to remove moss and lichens from the training dataset to improve the classification accuracy. We also removed shrubland, herbaceous wetlands, permanent water bodies and croplands due to their low representation across the region, as well as permanent snow due to the challenges in accurately classifying this land cover (Appendix: Figure A5). Thus, our reference data includes bare/sparse vegetation (barren), forest and grassland classes.

##### b. Supervised classification algorithm

To predict the most probable land cover class at each site for each time period, we used the Classification and Regression Trees (CART) algorithm (Breiman et al., 2017). CART is a machine-learning algorithm that uses decision trees to predict the likelihood of a target variable falling in a specific class. The goal was for the CART algorithm to predict which spectral signature from Landsat corresponds best to the input training dataset, with each internal node limiting the loss function (i.e., the difference between the model’s predictions and the data). Every pixel was selected based on its surface reflectance value for that specific band and sent to the subsequent attribute and threshold decision node. We used the Recursive Partitioning and Regression Trees (rpart) package in R (v.4.4.2) to run the decision trees.

##### c. Training dataset

First, we merged the reference datasets from each study area into a regional dataset (n = 57315 pixels). We masked the pixels belonging to land cover classes other than forest, grassland and barren out of the rasters to limit errors during the supervised classification. To best calibrate the classifier, we balanced the training dataset by selecting the number of pixels found in the least represented land cover class (n = 3454) and randomly sampled the same number of pixels in the other, more represented classes. Then, we randomly sampled 30% of the balanced dataset (n_30%_ = 3108). From the sampled sites, we extracted surface reflectance values from both 2021 Landsat multitemporal metric composite, the pseudo-natural color composite (SWIR1, NIR and red) and the double shortwave composite (NIR, SWIR1, SWIR2). Thus, the two obtained input datasets included a sample of pixel coordinates, the associated 2021 composite reflectance values (i.e. NIR, SWIR1, and red or NIR, SWIR1, SWIR2), and the 2021 ESA land covers. The algorithm subset each input dataset and repeatedly classified them until every pixel belonged to a class. We selected 200 observations as the minimum number required in a node to generate a split (Appendix: Figure A6). Once trained, we used the classifier to classify the entire Landsat annual median composite, which assigned a probability of the likeliest land cover class for each pixel (Appendix: Figure A7). We selected the land cover with the maximum probability at each site and compared the predicted classification with the observed ESA classification (Appendix: Table A3).

##### d. Classification accuracy estimation

To evaluate the algorithm’s accuracy, we calculated the overall accuracy (Alberg et al., 2004), defined as the probability of an individual pixel being correctly classified, as well as the user’s accuracy (the number of commissions), the producer’s accuracy (the number of omissions) and Cohen’s kappa (κ), the measure of agreement between variables (McHugh, 2012) (Appendix: Table A4). We used k-fold cross-validation, randomly dividing the set of observations into k-folds (k = 5) of equal sizes to estimate the accuracy. We used the algorithm to classify Landsat mosaics for the remaining time periods (i.e., the five-year spans ending in 2001, 2005, 2009, 2013 and 2017), selecting the land cover with the maximum probability at each site to detect changes in land cover between time periods.

### 2.4. Analyses

#### 2.4.1. The role of topography in current land cover distribution

To model the role of topography on the distribution of land cover types across space, we used boosted regression trees (BRTs), a machine-learning framework using the XGBoost 1.7.8.1 package (Chen & Guestrin, 2016) in R (v.4.4.1). Boosted regression trees use a boosting algorithm and can handle a variety of data types, including non-linear data, as well as missing data and non-collinear data. By fitting many decision trees, BRTs can handle complex interactions while minimizing the loss function (Elith et al., 2008), also known as the error function. We used the results of our land cover supervised classification for each area for 2021.

For each pixel (n = 3105), we included the elevation, slope (degrees), northness, eastness and radial ruggedness index (RRI) generated during the topography extraction. Pairwise correlations among all variables indicated some correlation between slope and elevation (r = 0.59), slope and ruggedness (r = 0.48), elevation and ruggedness (r = 0.41) (Appendix: Figure A8). Correlations among the remaining variable ranged from -0.08 to 0.07. All correlations were low enough to not be a concern for this analysis. Using the classified satellite composites, we assigned a presence-absence binary value for each land cover class at each pixel (i.e., barren, grassland and forest).

We ran individual BRT analyses for each class using a logistic regression. The first step in running a BRT requires setting hyperparameters to minimize the root mean squared error (Appendix: Figure A9, Table A5 and A7). These parameters include the learning rate (eta), maximum loss reduction (gamma), the maximum tree depth (max_depth), and the subsample ratio of the training sample (sub_sample). We bootstrapped the model using different combinations of hyperparameters, resulting in 81 iterations, and extracted their respective root mean square errors. We then selected the set of hyperparameters that produced the lowest root mean square error (Appendix: Table A5).

To fit the final model, we split the dataset in an 80-20% training-test set, randomized the data and bootstrapped the models 1000 times (e.g. Sherman et al., 2023; Yan et al., 2021). We calculated each variable’s relative importance and effect on the land cover classes occurrence (Appendix: Table A8). In addition, we subtracted the predicted probability of land cover occurrence by the input binary classification (i.e., land cover with maximum probability at a site = 1, other classes assigned 0) to calculate bias. If the measures are close to 0, the model minimizes the prediction bias (Appendix: Table A6).

#### 2.4.2. Detection of land cover change

##### a. Quantifying change

To quantify the impacts of the sudden decrease of livestock abundance in the post-Soviet crash in the 1990s and its drastic increase by the 2020s, we used the 2001 and 2021 multitemporal Landsat composites generated from the supervised classification to quantify the overall change in each land cover surface area between the two endpoints of the time series.

Using the raw data, we summarized the number of pixels found in each land cover class and calculated the differences in surface area and overall change between 2001 and 2021. We repeated the same steps including the study area to understand where changes might be occurring in the region with the expectation to observe a decrease in grassland cover as pressure intensities increase.

##### b. Detecting the direction of change

To detect the direction of change, we selected pixels that experienced a change in land cover at any moment between 2001 and 2021 by using the classified multitemporal Landsat composites for the 6 time periods (i.e. 2001, 2005, 2009, 2013, 2017, 2021). We summarized the number of pixels changed between each period, by study area, elevation band (n = 5) as well as the main land cover transitions (Appendix: Table A9).

To quantify the direction of change in absolute terms, we scored whether the pixel changed in land cover (value = 1) or not (value = 0) between 2001 and 2021. We then binned the pixels by elevation, period, and disturbance intensity, and recorded the total number of pixels in each bin, as well as the number of pixels that had changed in either direction (i.e., from grassland to barren: “browning”; from barren to grassland: “greening”) (Appendix: Table A10).

To determine the role of elevation, time and disturbance on the proportion of land cover change, we ran a generalized linear mixed-effects model with a negative binomial distribution and log link function using the glmmTMB package (Brooks et al., 2017). We modeled the number of pixels that changed land cover type as a function of pressure intensity (i.e., study area), elevation, period, and land cover transition type (i.e., browning or greening; forestation was omitted because of the low sample size), as well as select two-way interactions. We used the logged total number of pixels in each bin as an offset term to allow model outputs to be interpreted as effects on the *proportion* of pixels that transitioned rather than the *number* (Appendix: Figure A11). We ran a single full model instead of performing model selection, as our sample size was sufficient to include all parameters of interest, and each parameter was associated with a clear hypothesis (Appendix: Table A11). We evaluated the model fit using the DHARMa package (Hartig, 2016), and the *emmeans* packages (Lenth, 2017) to perform post-hoc pairwise comparisons between transition types. We also calculated the estimated marginal mean for each bin using a Tukey adjustment for multiple comparisons.

## 3. Results

### 3.1. Land cover classification

The overall accuracy of the supervised classification was high (OA = 0.86), with a substantial agreement (κ = 0.78), high probability of accurately classified pixels (PA = 0.86), and classification reliability (UA = 0.86). The reliability of the classification was the lowest for grasslands (UA = 0.76; Appendix: Table A4).

#### 3.1.1. Determinants of current land cover distribution

The relative importance of each topographic variable in predicting the 2021 distribution of land cover classes varied per class. Elevation best predicted the distribution of barren areas (RI_mean_ = 0.82), followed by slope, ruggedness, and eastness (RI_mean_< 0.1) (Figure 3, Table A8). All topographic variables played a similar role in predicting forest land cover, with a higher relative importance of elevation (RI_mean_ = 0.44) and northness (RI_mean_ = 0.24) than slope (RI_mean_ = 0.12), eastness (RI_mean_ = 0.11) and RRI (RI_mean_ = 0.08). Elevation remained the most important predictor for grasslands (RI_mean_ = 0.64), followed by northness (RI_mean_ = 0.14), slope (RI_mean_ = 0.08), eastness and RRI (RI_mean_ = 0.07). Overall, the best topographic predictor for land cover distribution was elevation. Forested areas primarily characterize lower elevations, while grasslands dominate mid-elevation, and barren areas constitute the main land cover type at high elevations, though they are also present at lower elevations but to a lesser extent (Figure 3b. B). Northness and eastness were the second-best predictors of current land cover distribution, with a greater proportion of forested areas located on northern slopes compared to grasslands, which are mostly found on southern slopes (Figure 3b. A-C). Ruggedness and slope did not contribute to explaining the distribution of any of the land cover classes (Figure 3b. D-E) (Appendix: Table A8).

**Figure 3.**
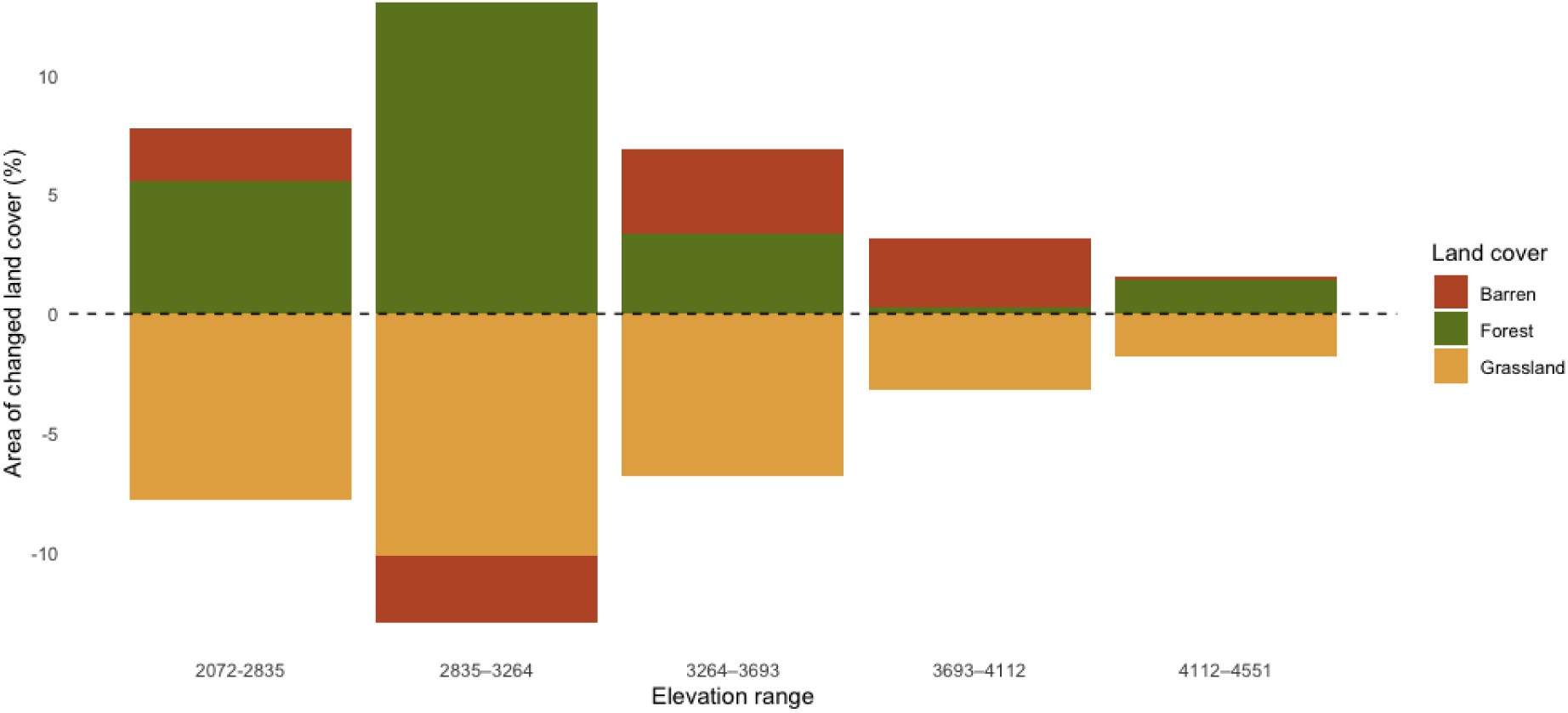
Proportional differences in cover of forests, bare/sparse vegetation and grasslands across elevation bands between 2001 and 2021. The proportion of changed land cover was calculated using the total count of pixels for each land cover class in each elevation band for 2001 and 2021. The difference in number of pixels between 2001 and 2021 was used to determine the area of change. Areal change for each land cover and study area was then divided by the total number of pixels in each band (n_band1_= 9370, n_band2_= 15769, n_band3_= 18331, n_band4_= 11808, n_band5_= 1531) and converted to percentages. The elevation bands include the elevation ranges across study areas and are organized from the lowest (1) to the highest (5). Positive values indicate increases in area over the study period.

### 3.2. Detection and direction of land cover change over time

#### 3.2.1. How much has changed overall?

Across all study areas and land cover types, 84.8% of pixels maintained the same land cover class between 2001 and 2021 (Appendix: Table A9). We found that grasslands remained the dominant land cover type across the landscape, covering 62.7% of the area, followed by barren (32.6%) and forest (4.7%) types in 2021(Appendix: Figure A10).

Of all pixels that changed between 2001 and 2021, grasslands experienced most change (9.1% of grassland pixels changed), followed by barren areas (4.7%). The proportion of forest cover that changed to another land cover type was less than one percent. Thus, we report the direction of change only for grasslands and barren areas. Of the 9.1% of grassland pixels that changed, the majority (67.7%) transitioned to barren and fewer (32.3%) to forest. Of the 5.4% of barren pixels that changed, 63% transitioned to forests and 37% to grasslands (Appendix: Table A10). Overall, these figures represent a 339-hectare loss of grassland area, a 274-hectare gain in forest area and a 65-hectare gain in barren area between 2001 and 2021, which were distributed unevenly across elevations and the gradient of anthropogenic pressures.

#### 3.2.2. Land cover change in relation to elevation

Grassland area was lost between 2001 and 2021 across all elevation bands, but the largest area in absolute terms was lost at low-intermediate (144 ha loss) and intermediate elevations (113 ha loss; Figure 4). The increase of forested areas mainly occurred at elevations ranging between 2835 m and 3264 m (185 ha gain; elevation band 2), 3264 m and 3693 m (55 ha gain; elevation band 3) and 2072 m to 2835 m (47 ha gain; elevation band 1) (Figure 4). We also observed an increase in barren areas across all elevation bands, with the exception of intermediate elevations where 39 ha of barren areas were loss.

**Figure 4.**
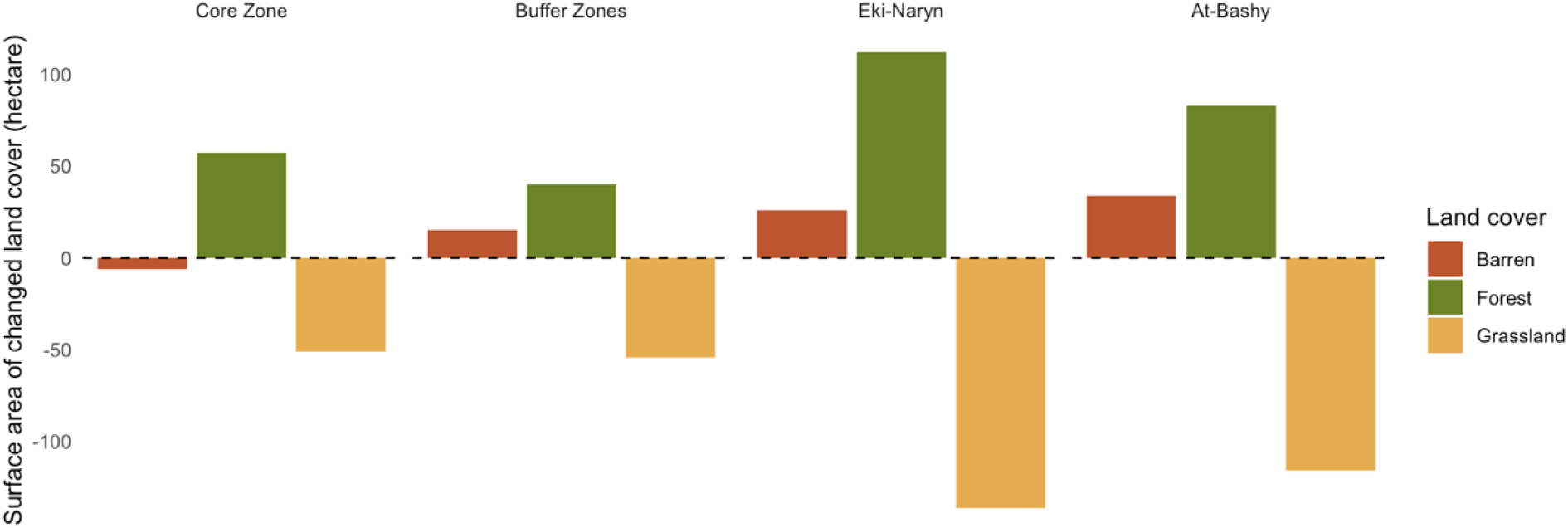
Change in spatial coverage of forested, barren and grassland areas between 2001 and 2021 in each study area. The proportion of changed land cover was calculated using the total count of pixels for each land cover class in each study area for 2001 and 2021. The area of change for each land cover and study area was then divided by the total number of pixels in each study area (n_core_ = 7997, n_buffer_ = 14647, n_eki-naryn_= 23118, n_at_bashy_ = 23146) and converted to hectares.

#### 3.2.3. Land cover change in relation to disturbance

All study areas were characterized by a net decrease in grasslands and increases in forested and barren areas between 2001 and 2021 (Figure 5). These changes appeared to co-vary with increasing anthropogenic pressures. The least disturbed site, i.e. the protected core zone of Naryn State Reserve, posted the smallest decline in grassland cover, which was 2-3 times less in absolute terms than the declines occurring at the two most disturbed sites (Figure 5). The patterns of increase in forest and barren cover mirrored each other, being lowest in the Naryn State Reserve and 2-4 times higher in areas with intermediate and high anthropogenic pressures (Figure 5).

**Figure 5.**
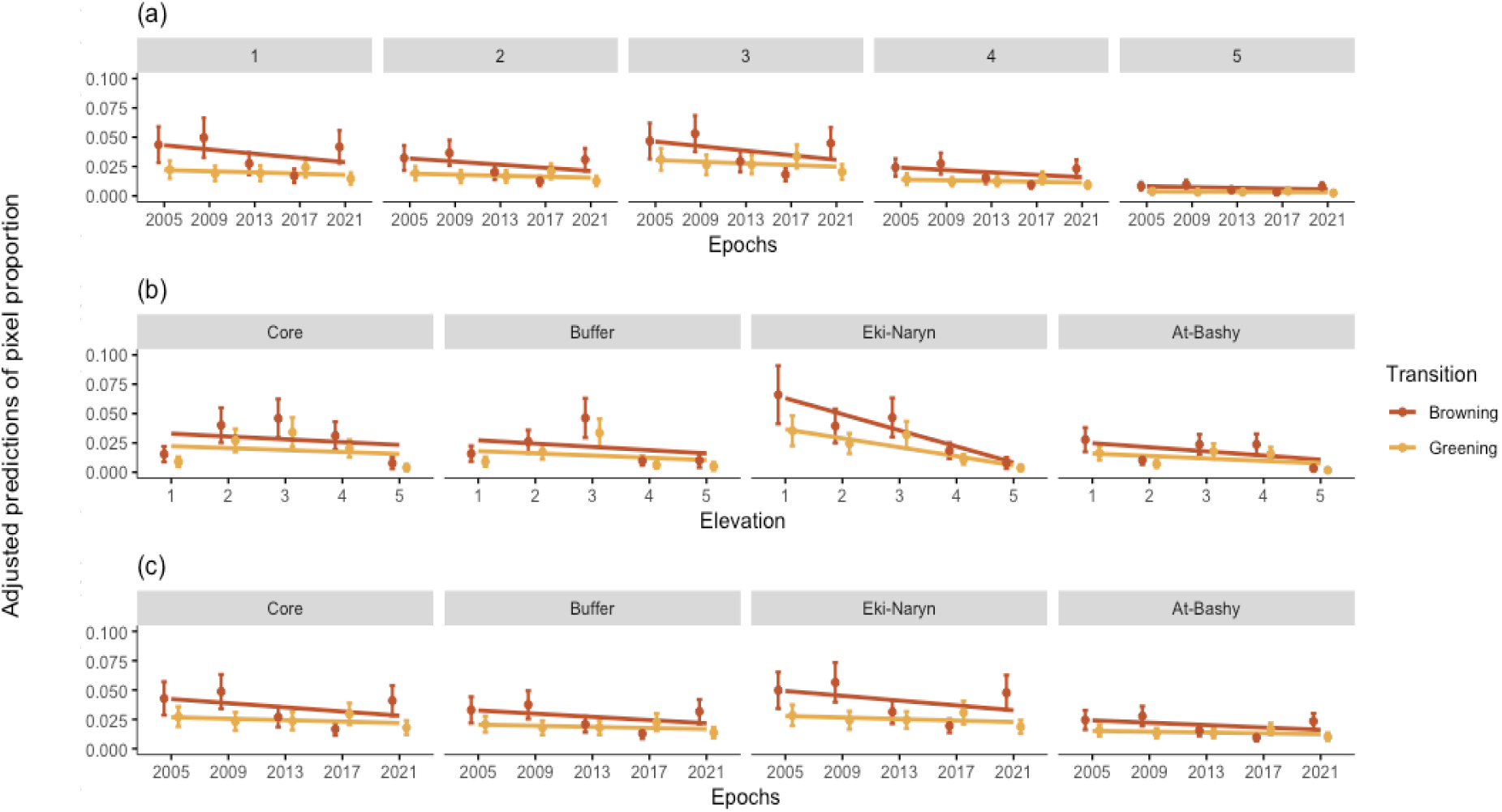
Adjusted mean predicted proportions of pixels that transitioned from grassland to barren (browning) and barren to grassland (greening). The transition types are reported in (a) each elevation band at each given time period ,(b) each elevation band in each area, and (c) each study area at each given time period. The study areas are ordered along a gradient of anthropogenic pressures, with the least disturbed area on the left (core) to the most disturbed area on the right (At-Bashy). Points represent estimated marginal means based on the model, and error bars represent 95% confidence intervals. Linear regression lines are plotted to better visualize trends.

#### 3.2.4. Interacting factors determining trends in land cover change

Due to the large number of interactions included in the model, individual model coefficients are challenging to interpret and are only presented in Appendix: Table A11. To facilitate the interpretation of the results, we ran post-hoc pairwise comparisons (using a Tukey adjustment for multiple comparisons) to answer biologically meaningful questions and present the results of these tests here. For each comparison, we indicate which levels of which variable are being compared. Unless otherwise indicated, the estimated effects for each level were marginalized over the levels of the other variables (e.g., estimated effects for different areas are averaged across years and elevations).

In general, more pixels transitioned to barren than to grasslands. Regardless of protection status, all areas experienced significantly more browning than greening when averaged across years and elevations (all pairwise contrasts between amount of browning and greening p < 0.05; Table 1).

There was no consistent pattern in the relative amount of browning and greening over time. In the five-year periods starting in 2005, 2009, and 2021, there were significantly higher proportions of browning than greening (p < 0.01, p < 0.001, p < 0.001, respectively), while in the period starting in 2017 there was a shift to significantly more greening than browning (p < 0.05; Table 1). There was no significant difference in proportions of browning vs greening in the period starting in 2013 (Table 1). The relative differences in the amount of browning and greening were driven largely because of inter-period changes in the proportion of browning. The proportions of greening stayed relatively constant across periods in most elevations and areas (Figure 6).

Across most elevations, there was significantly more browning than greening (Table 1). However, there was a lot of variability in elevation patterns based on the area (Figure 6). For instance, in the core and buffer zones, the proportion of pixels changed peaked at intermediate elevations (pairwise contrasts between elevation band 3 and elevation bands 1 and 5 for both areas: p < 0.05; Appendix: Table A14; Figure 6). In Eki-Naryn, the proportion of change was highest at low elevations (pairwise contrasts between elevation band 1 significantly different than elevation bands 4 and 5: p < 0.001; Appendix: Table A14; Figure 6). At-Bashy had the lowest overall proportion of change and no consistent pattern across elevation.

## 4. Discussion

With unprecedented pressures from anthropogenic activities and climate change directly challenging the productivity and dynamics of mountain habitats, it is crucial to understand localized responses to better inform local management. Yet, we are often limited by the large spatiotemporal resolutions required to characterize the responses. Here, we identified intermediate elevations, high degree of disturbances, and the interaction between the two as the main contributors of recent change in the cover of grasslands, forested and barren habitats, ultimately resulting in a net loss of grassland and increases in forests and barren areas. These findings were revealed by combining Landsat analysis-ready data (ARD) and the European Space Agency (ESA) world cover dataset in a way that is applicable to many data-poor, remote regions of the world.

### 4.1. Role of topography in current land cover distribution

Topography plays a crucial role in the distribution of terrestrial ecosystems (Chiu et al., 2020). We found elevation to be the most important predictor of forest, grassland and barren land cover types, which is consistent with the previously established relationships between elevation and thermal gradients that drive species’ niche distribution in mountain ecosystems (Chauvier et al., 2021; Fang et al., 2024; Guo et al., 2018). Indeed, the increase in barren and sparse vegetation surface area as elevation increases directly links to abiotic conditions that limit the establishment of plant communities at altitudes beyond their physiological tolerance range (Körner & Diemer, 1987). While our results also suggest that elevation may explain the occurrence of barren areas at low elevations, we later explore the role anthropogenic pressures play in land cover distribution.

Aspect was also an important determinant of current grassland and forest distribution. Grasslands were mostly found on south-facing slopes with minimal terrain heterogeneity, while forested sites characterized north-facing slopes. Southern slopes are more exposed to solar radiation, providing grassland communities with greater opportunities to grow owing to warmer temperatures and prolonged foraging opportunities for herbivores. Warmer temperature also results in faster and complete snow melt, which improves the release of water at the beginning of the growing period and leads to the full exposure of the landscape to solar radiation throughout the short growing season. In contrast, northern-facing slopes are less exposed to solar radiation, which leads to less extreme warm temperature and a longer establishment of snowpack that gradually melts through the year. The presence of snowpack provides shaded and moist microhabitats ideal for alpine tree and shrub species with less tolerance to drought (Måren et al., 2015; Verrall et al., 2023), leading to localized increased productivity and thermoregulation opportunities for wildlife.

While ruggedness and slope did not explain the distribution of grasslands, forests and barren areas in our study region, a more complex terrain heterogeneity has been shown to result in diverse topo climates, leading to niche differentiations and the establishment of diverse and highly specialized alpine communities (Antonelli et al., 2018). The evolution of plant traits adapted to a specific set of biotic and abiotic conditions increases the sensitivity of these communities to shifts in environmental conditions and anthropogenic pressures (Adler et al., 2014; Borchardt et al., 2013; Meineri et al., 2020). Thus, the inclusion of terrain heterogeneity and complexity remains important when disentangling the drivers of land cover distributions at more localized spatial scales and to better identify which parts of the landscape is more sensitive to shifts in climatic patterns and increased anthropogenic pressures. Using more accurate land cover classifications and topographic information at finer spatial resolutions may reveal patterns not detected at the current scale.

### 4.2. Magnitude of land cover changes over two decades

Perhaps our most surprising finding was that most pixels (∼84%) retained the same dominant land cover between 2001 and 2021. This is contradictory to our prediction following the extensive report of pasture degradation in the area over the last decade (Domenech et al., 2022). The absence of field records limited our ability to evaluate whether grasslands were undergoing intra-state degradation, as highlighted in studies focusing on weed encroachment and the increase of unpalatable species. As such, these results are mostly indicative of a methodological bias.

However, a change of ∼15 % in land cover types over 20 years in an arid ecosystem can still be considered important. A small proportion of pixels that changed land cover type translated into an overall loss in grassland area and gains in forest and barren area. These losses and gains are modest in absolute terms: 339 ha of grassland were lost, while 65 ha of barren area and 274 ha of forest were gained. In general, more grassland was lost, and conversely forest and barren area gained, at lower to middle elevations (i.e. bands 1-3, < ∼3,700 m), and along the gradient of increasing disturbance. These findings are consistent with the grassland degradation and desertification (Bardgett et al., 2021; Yan et al., 2023; M. Zhang et al., 2025) and the increase of forested areas observed worldwide (Robinov et al., 2021; Stockdale et al., 2019; L. Zhang et al., 2022) in montane regions.

A net increase in forested area is of interest because where it is occurring. The largest increases in forested areas occurred between 2835 m and 3693 m (i.e., bands 2 and 3), which coincides with the upper range limit of Schenk’s spruce (3500 m; Zhang et al., 2016), the dominant species in Naryn forests. The expansion of Schrenk spruce within and beyond their known upper range limit suggests that warmer temperature may play a role in driving the upward shifts of treelines in the region, as reported in other alpine ecosystems (Dullinger et al., 2004; Hansson et al., 2021; Harsch et al., 2009; Qi et al., 2015; Wang et al., 2006). The sensitivity of Schrenk spruce to warmer temperature supports the hypothesis that higher elevations may be a climate refuge for the species (Qin et al., 2020, 2022; R. Zhang et al., 2016). Despite forest encroachment being considered negative in the context of grassland conservation, the increase of forested areas can be beneficial by providing new habitats for the local biodiversity and ecosystem services, such as forestry. The increase of forested areas mainly occurred on northern slopes where grazing by livestock was less important. This result highlights the role of grazing in limiting forest encroachment, and therefore grassland contraction in some areas.

### 4.3. Direction of land cover changes over two decades

Pixels that transitioned did so in a relatively predictable way, with most transitions occurring between grasslands and barren habitat types (browning vs greening). Although there was no significant effect of time, disturbance, or elevation, we observed a net increase of barren habitats across all study areas, at the exception of the highly protected core zone, which was the only area with a decrease in barren areas. The core zone was also characterized by the lowest loss of grasslands amongst all study areas, which indicates that, in the absence of direct anthropogenic pressures, grasslands may decline in response to the warming trends in the country (Guo et al., 2018). Indeed, by including the core zone of the reserve, which does not include any livestock since 1983, we can better compare response to changes across study areas, and understand which changes are driven by grazing and/or shifts in thermal and precipitation regimes. Mild disturbances in the buffer zones yielded to similar loss of grasslands as in the protected core zone but the increase of barren areas indicate that even mild anthropogenic pressures may result in habitat degradation. Moreover, we found compelling evidence that increasing anthropogenic pressures lead to substantial browning, potentially amplifying already existing climate-driven impacts (Y. Yang et al., 2023). However, and to our surprise, we found that areas with prolonged exposure to high degrees of anthropogenic disturbances, such as At-Bashy, were characterized by a lesser magnitude of land cover change and browning compared to Eki-Naryn, which is affected by intermediate levels of disturbances. Our observations suggest that prolonged exposure to high anthropogenic pressures continues to negatively impact grassland habitats, while promoting the expansion of barren and forested area, but less than in areas characterized by lower disturbance pressure. A comparison with recent records of pasture degradation in At-Bashy (Domenech et al., 2022) raises the question of whether the observed patterns align with preliminary signs of a regime shift within grasslands (Y. Zhao et al., 2023).

### 4.4. Strengths and limitations of our approach

Changes in alpine plant communities in response to anthropogenic and climate-induced stressors tend to be subtle and gradual. Detecting and characterizing these changes require long-term and frequent observations at fine spatial resolutions. Our methods showcase the potential of using GLAD Landsat ARD (Potapov et al., 2020) to map and characterize localized land cover distribution and change since the late 1990s. The production of localized mapping and characterization of land cover distribution are particularly important in areas with limited records of plant traits and diversity. Although temporal gaps in Landsat imagery remain the main limitation in areas such as Naryn due to cloud cover and sensor failure (Roy et al., 2016), the GLAD Tools address this issue through gap-filling methods to improve data quality by generating cloud-free, multi-annual Landsat composites. Multi-annual composites can then be used to classify land cover over time, which we did retrospectively using a set of training data from ESA’s 2021 land cover (Zanaga et al., 2022). In the absence of ground-truthing data (i.e., vegetation records at recorded locations, high resolution imagery) characterizing local land cover types and distribution in 2021, the 10 m resolution of the ESA WorldCover provided the most accurate spatial characterization of land cover distribution across the landscape. Combining ESA imagery, Landsat multi-annual composites and SRTM digital elevation models enabled us to establish a baseline characterization of topographic and land cover within each study area. This baseline then enabled us to dive deeper into the mechanisms and drivers of land cover change in alpine communities.

Nevertheless, there are some caveats to our approach. While we successfully identified the spatio-temporal distribution and variations in grasslands, forests and barren areas, the absence of site-specific environmental, species and community-level information limited our ability to locally validate land cover classification and detect mixed land cover types. To address this issue using remote sensing methods, an analysis including mixed pixels may drastically improve the detection of areas sensitive to change, and provide a more accurate depiction of the landscape dynamics over time and space (Hermosilla et al., 2022; C. Yang et al., 2017).

Moreover, obtaining local maps of the historical and contemporary distribution of pastures based on their seasonal use would help delineate elevational ranges of interest to better map and characterize land cover dynamics based on known seasonal usage of the landscape. When characterizing land cover types in alpine regions, we highlight the importance of recognizing the variation in sample sizes between high and mid-elevations. Indeed, high elevations are characterized by smaller surface areas compared to the intermediate and low elevations, which is directly reflected in sampling effort (Appendix: Table A2). The variation in surface area can drastically limit the detection, quantification and comparison of dynamics related to change and requires careful interpretation.

## 5. Conclusion

This study aimed to assess fine spatial-scale patterns of rangeland degradation using satellite imagery to classify, quantify and map land cover distribution over multiple decades, evaluating the influence of topography, anthropogenic disturbances and climate change (through proxies) on land cover dynamics. These objectives were successfully met through the development of a robust framework that identified elevation and northness as the primary topographic drivers of land cover distribution, along with the role of protected areas and mild anthropogenic disturbances in mitigating landscape degradation. While the combined used of harmonized Landsat ARD imagery, ESA global land cover maps and digital elevation models at fine spatial scales provided valuable contemporary and historical landscape and habitat-level insights, future work must integrate a more nuanced pixel characterization, as habitats are often mosaics of vegetation types and land uses with gradual changes that do not conform to discrete categories. Collecting field data, such as comprehensive surveys of plant diversity, information on seasonal pasture types, site-specific disturbances and local weather will also be essential to accurately disentangle and identify the drivers of land cover change.

## Author contributions

Sherry C.E. **Young**: Conceptualization, Methodology, Validation, Formal Analysis, Investigation, Resources, Data Curation, Writing – Original Draft. **Steven Brownlee**: Methodology; Formal Analysis; Data Curation; Writing – Review and Editing. **Helen F. Yan** and **Hannah Watkins**: Methodology; Formal Analysis; Data Curation; Writing – Review and Editing. **Isabelle M. Côté**: Supervision, Writing – Original Draft, Review & Editing.

## Acknowledgements

We would like to thank Dr. Cole Burton (University of British Columbia), Dr. Chelsea Little (Simon Fraser University), Dr. Jonathan Moore (Simon Fraser University), Dr. Ruth Joy (Simon Fraser University), Dr. Colleen Cassady St Clair (University of Alberta) and Dr. Ron Ydenberg for their valuable input on the manuscript.

## Supporting Information and Data Availability

Supporting Information is available as part of this submission. Code is available on <u>GitHub</u> and the data is accessible through Google Earth Engine and GLAD Tools.

