## Appendix for "Multidecadal changes in land cover across a disturbance gradient in mountain grasslands of Kyrgyzstan"

Running Headline: Change in alpine rangelands

Author(s), and Corresponding Author(s)*

Sherry C.E. Young*

Simon Fraser University, 8888 University Drive, Burnaby, BC, V5A 1S6, Canada

University of Alberta, 11011-88 Avenue, Edmonton, AB, T6G 2G5, Canada

ORCID: 0000-0001-9757-0517

Hannah V. Watkins
Simon Fraser University, 8888 University Drive, Burnaby, BC, V5A 1S6, Canada

Steven Brownlee

Simon Fraser University, 8888 University Drive, Burnaby, BC, V5A 1S6, Canada

Helen F. Yan

James Cook University, 1 James Cook Dr, Douglas QLD 4814, Australia

Isabelle M. Côté

Simon Fraser University, 8888 University Drive, Burnaby, BC, V5A 1S6, Canada

**Workflow**

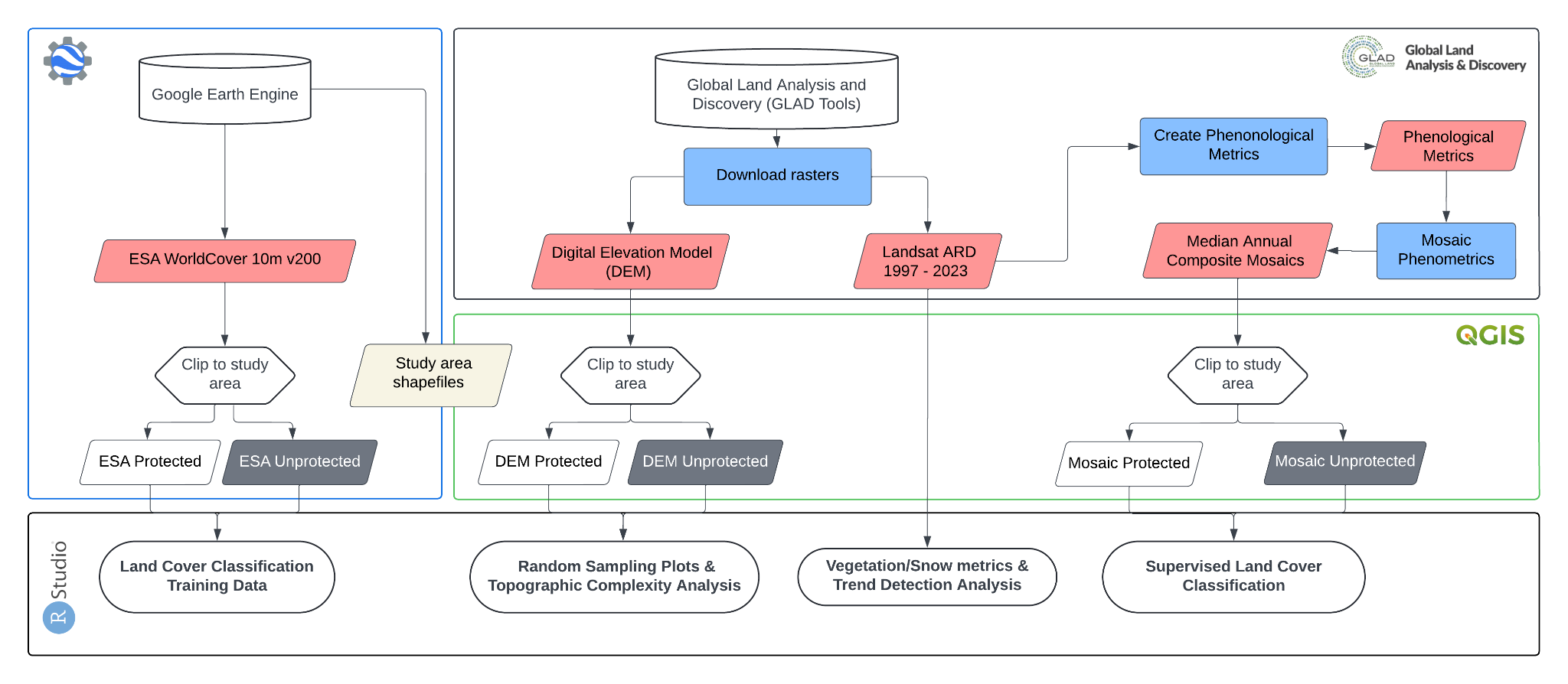

Figure A1.	Data sources and preparation flow chart. The GLAD Tools protocol enables the download of the digital elevation models (DEM) and Landsat ARD imagery.

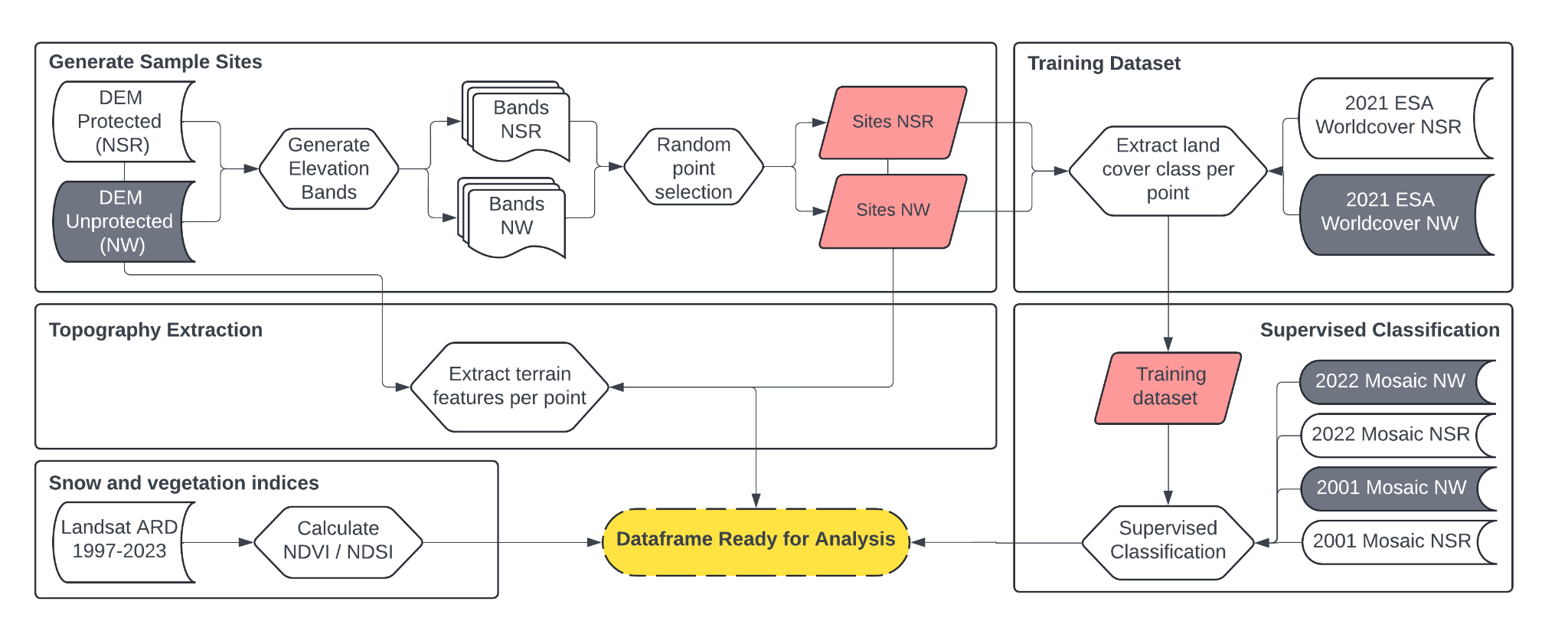

Figure A2.	Flowchart describing the data processing steps.

Study areas elevation band and point selection

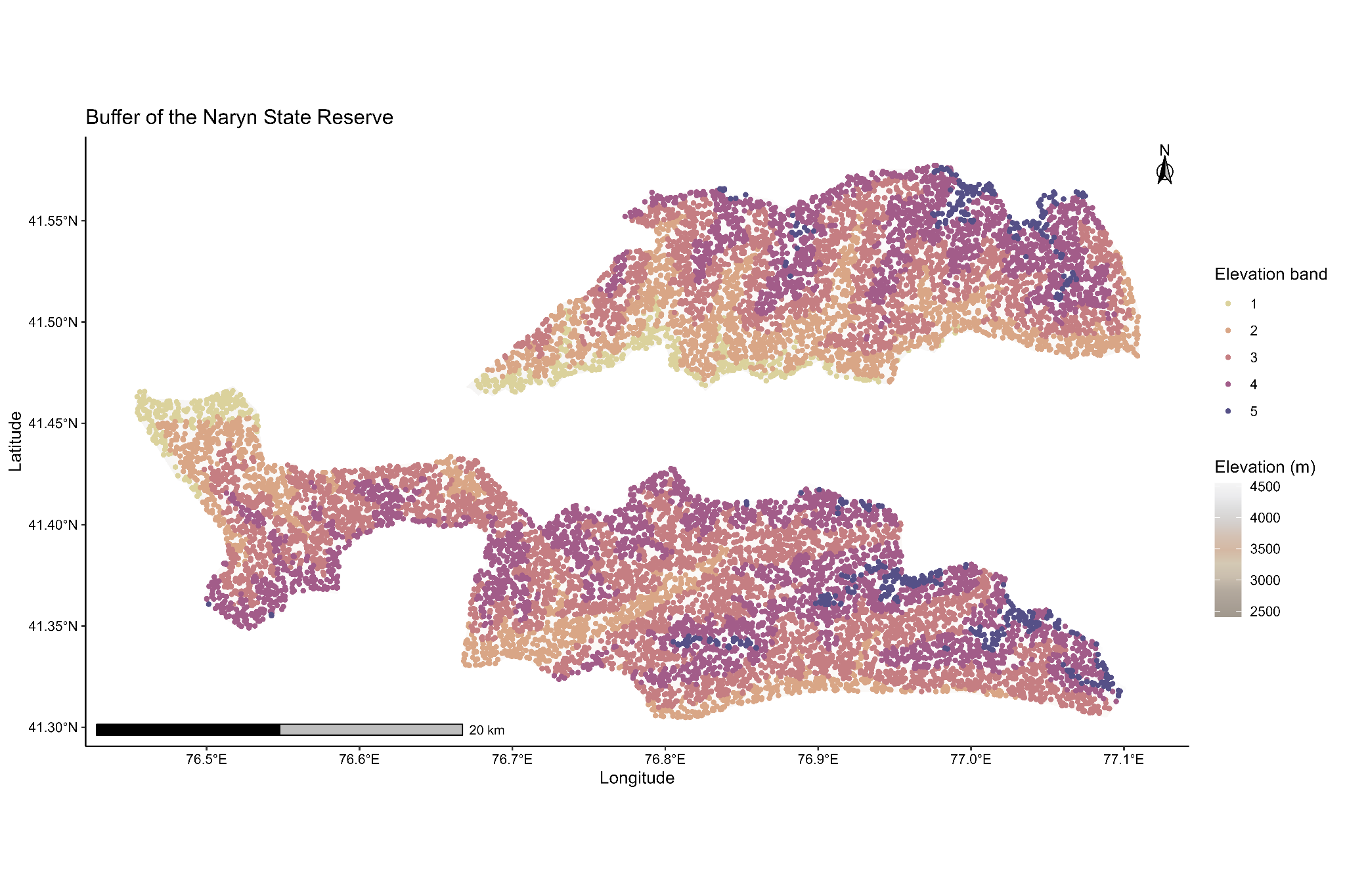

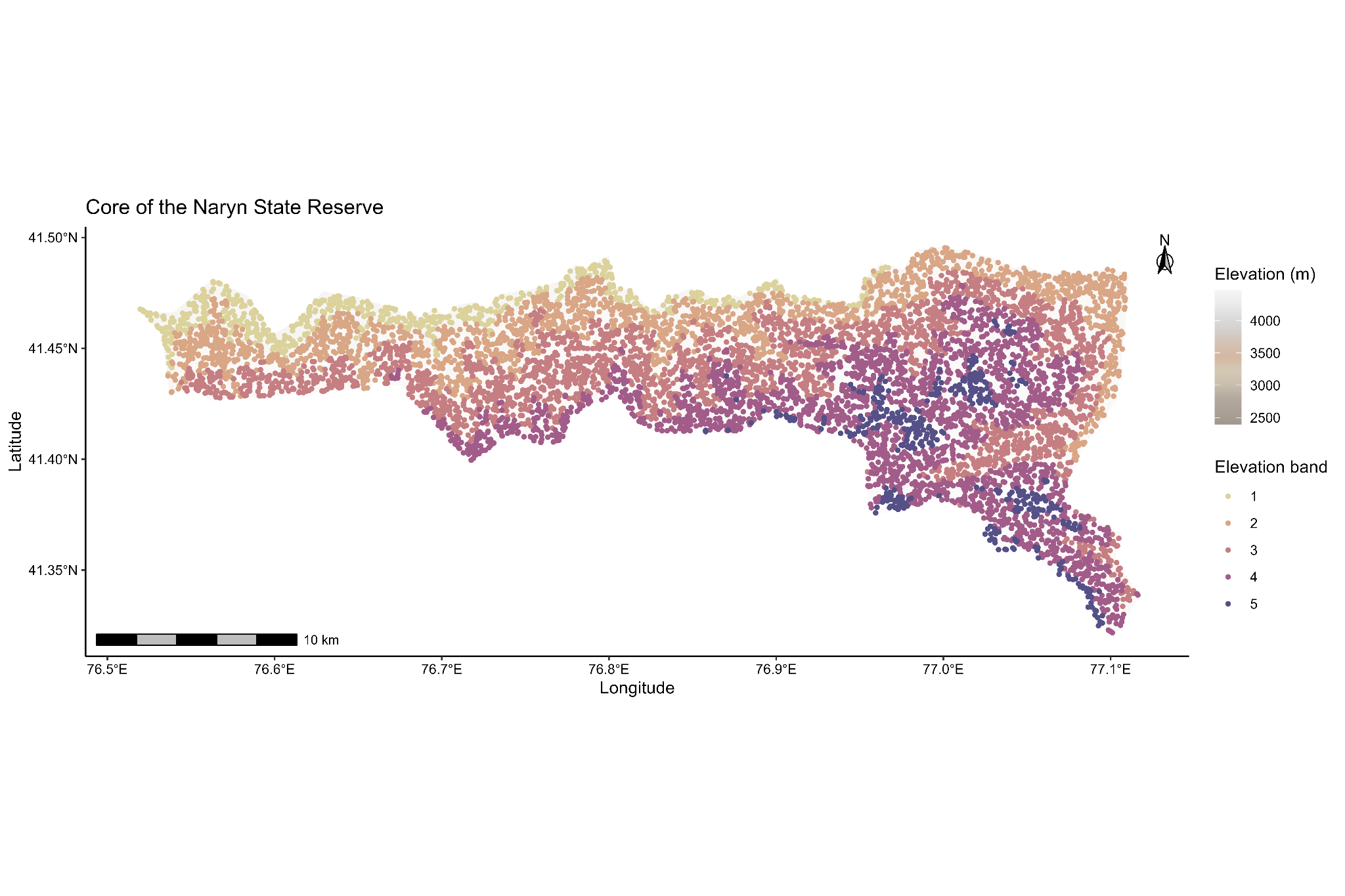

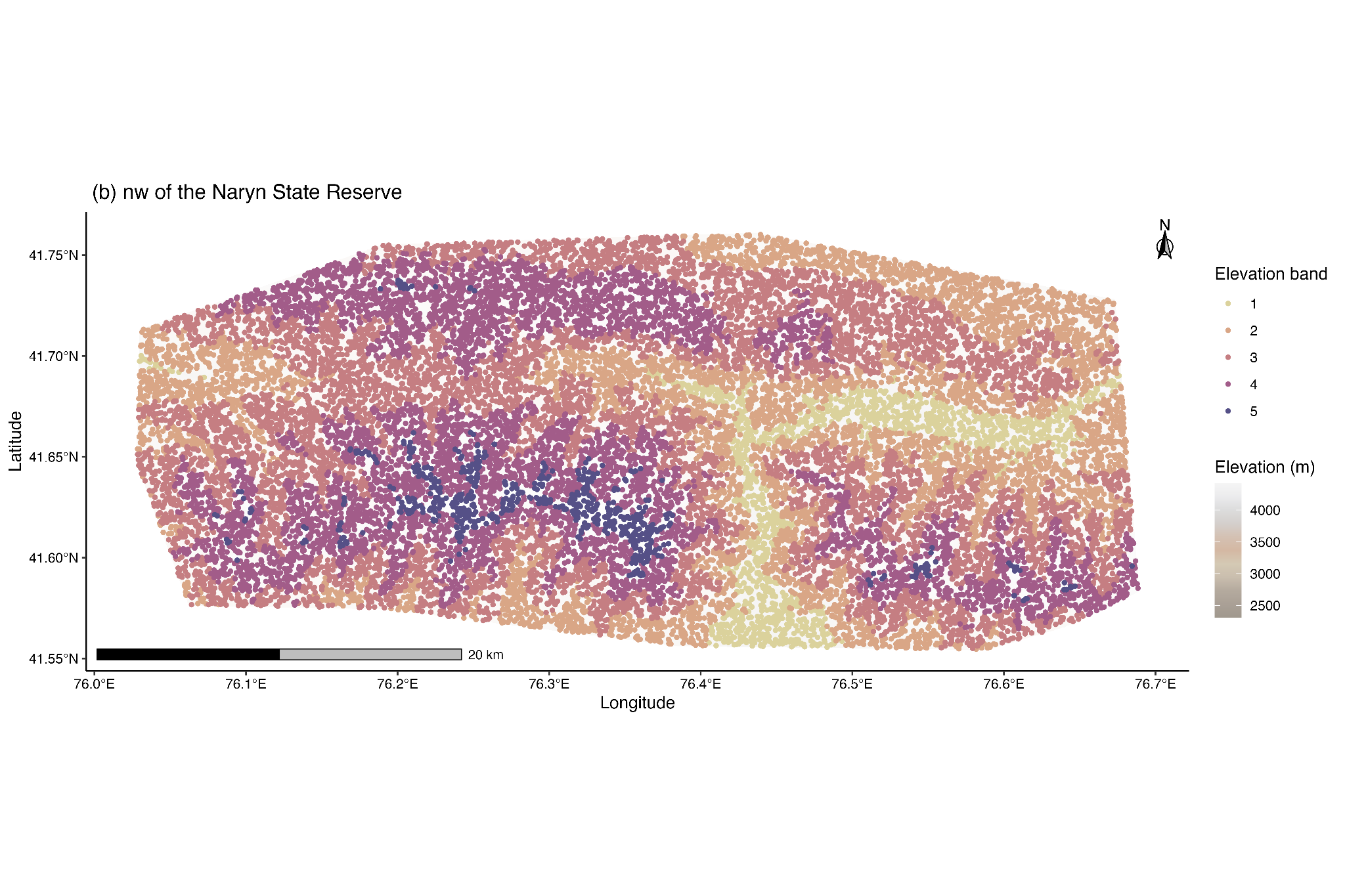

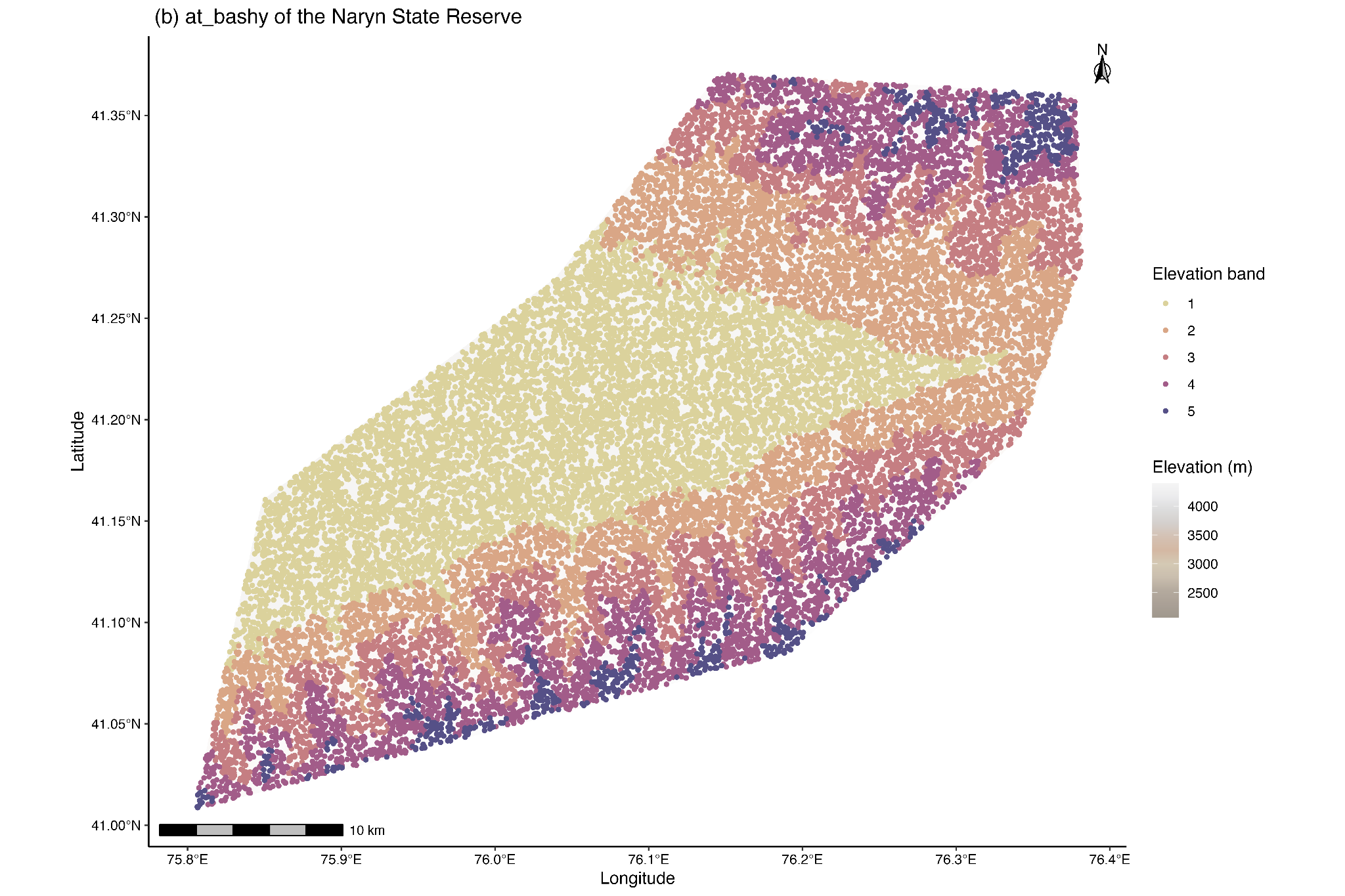

Figure A3.	Random selection of points for land cover analysis in the core zone of the Naryn State Reserve (top left) and the buffer zones (top right), the Eki-Naryn region (bottom left) and At-Bashy (bottom right) per elevation band (low = 1, high = 5). The darker colours indicate a higher elevation.

Table A1.	Elevation range (m) of each elevation bands for each study area

|  | Elevation bands | | | | |
| --- | --- | --- | --- | --- | --- |
| Area | 1 | 2 | 3 | 4 | 5 |
| Core zone | 2396 - 2811 | 2811 - 3225 | 3225 -3640 | 3640 - 4054 | 4054 - 4469 |
| Buffer zones | 2407 - 2835 | 2835 - 3264 | 3264 -3693 | 3693 - 4112 | 4112 - 4551 |
| Eki-Naryn | 2313 –2734 | 2734 –3156 | 3156 –3577 | 3577 –3999 | 3999 - 4420 |
| At-Bashy | 2072 - 2536 | 2536 - 3001 | 3002 -3466 | 3466 - 3931 | 3931 - 4396 |

Table A2.	Sample size of randomly selected pixels at elevation band per area using 1% of the total available sample size.

|  | Elevation bands | | | | |
| --- | --- | --- | --- | --- | --- |
| Area | 1 | 2 | 3 | 4 | 5 |
| Core zone | 638 | 1889 | 2605 | 2418 | 447 |
| Buffer zones | 618 | 2970 | 5999 | 4560 | 500 |
| Eki-Naryn | 1563 | 5733 | 8556 | 6572 | 694 |
| At-Bashy | 8064 | 5752 | 4272 | 4104 | 954 |

ESA Land cover classification

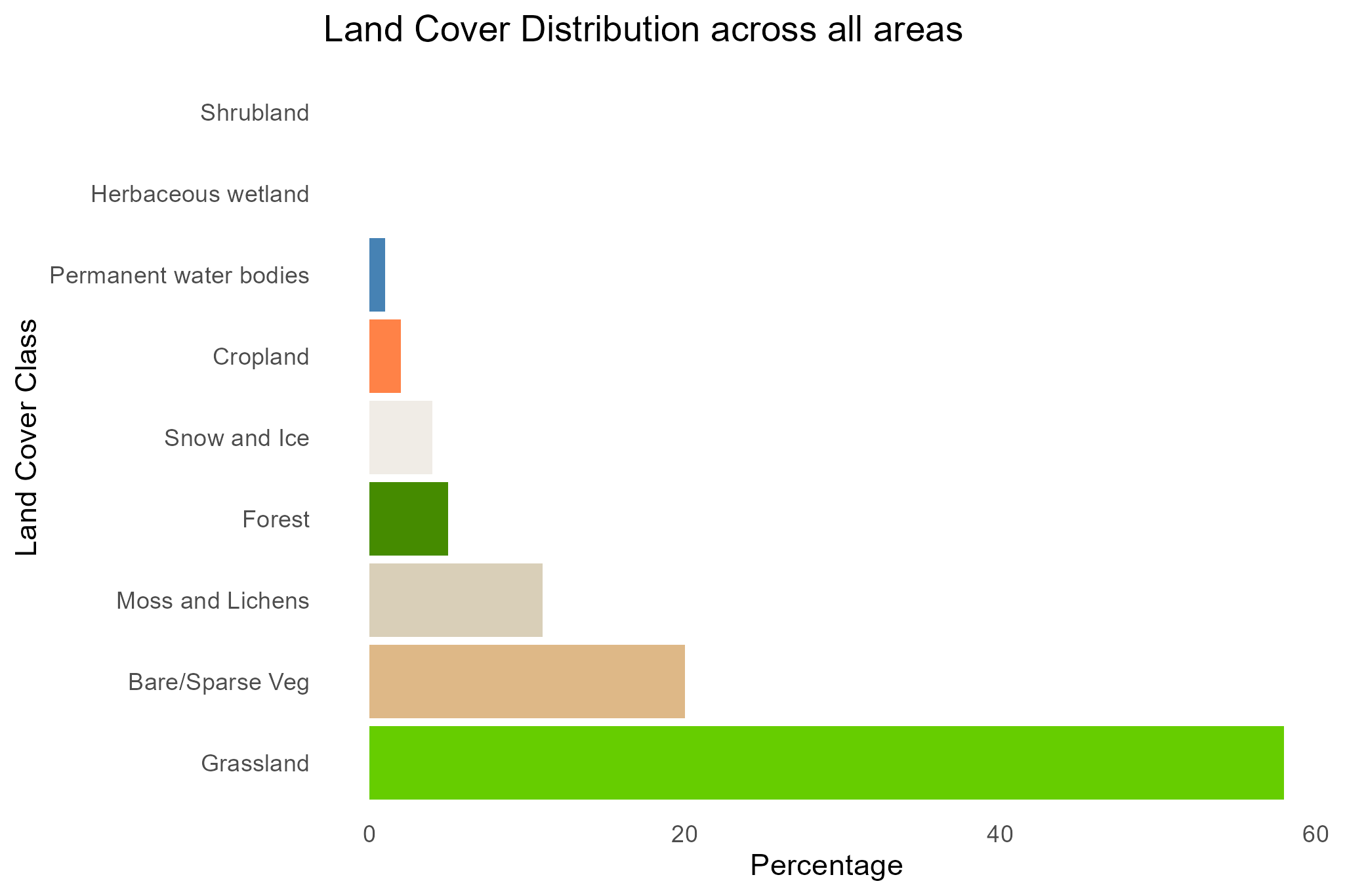

Figure A4.	Percentage of land cover class cover in the Naryn region.

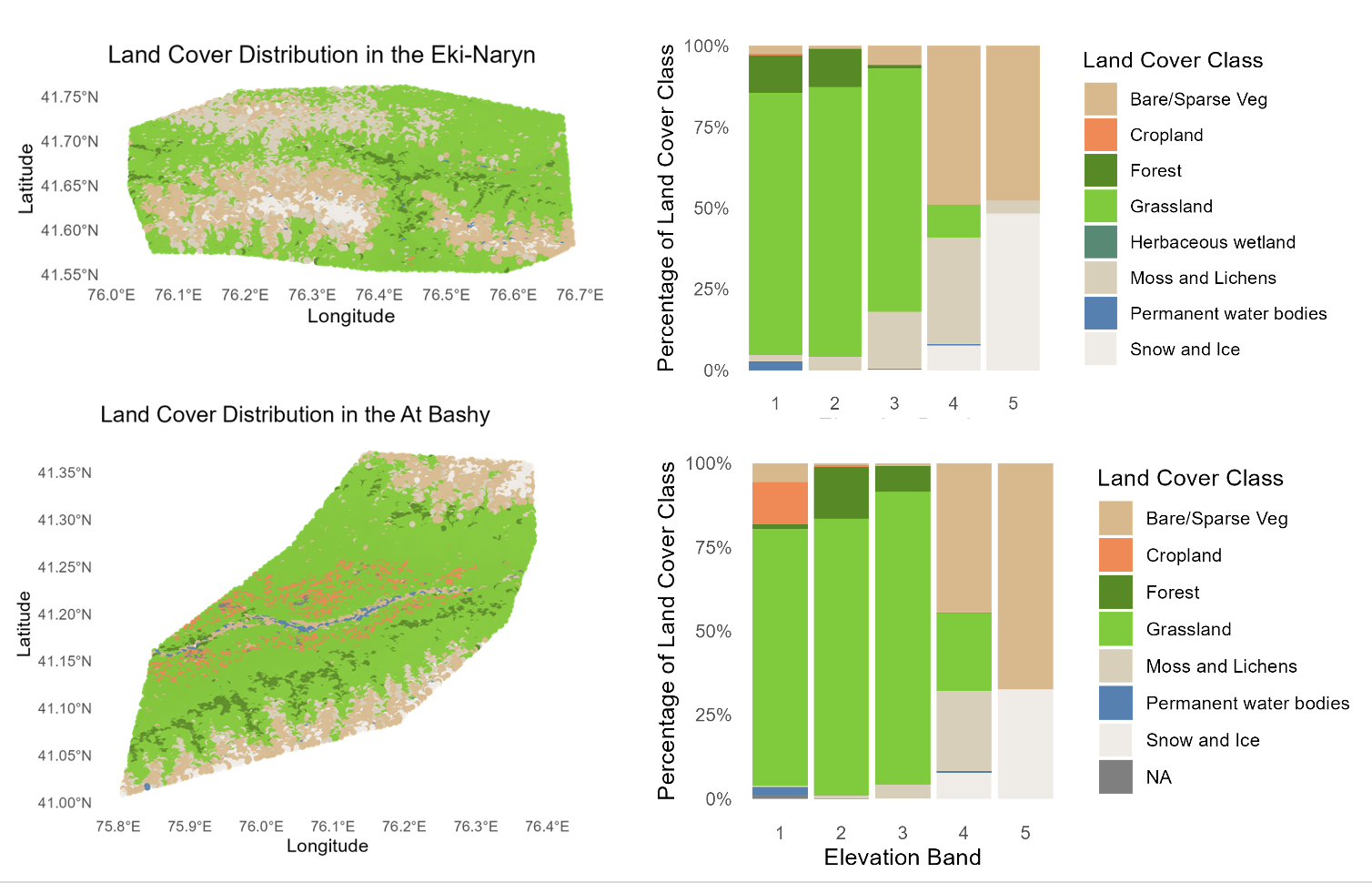

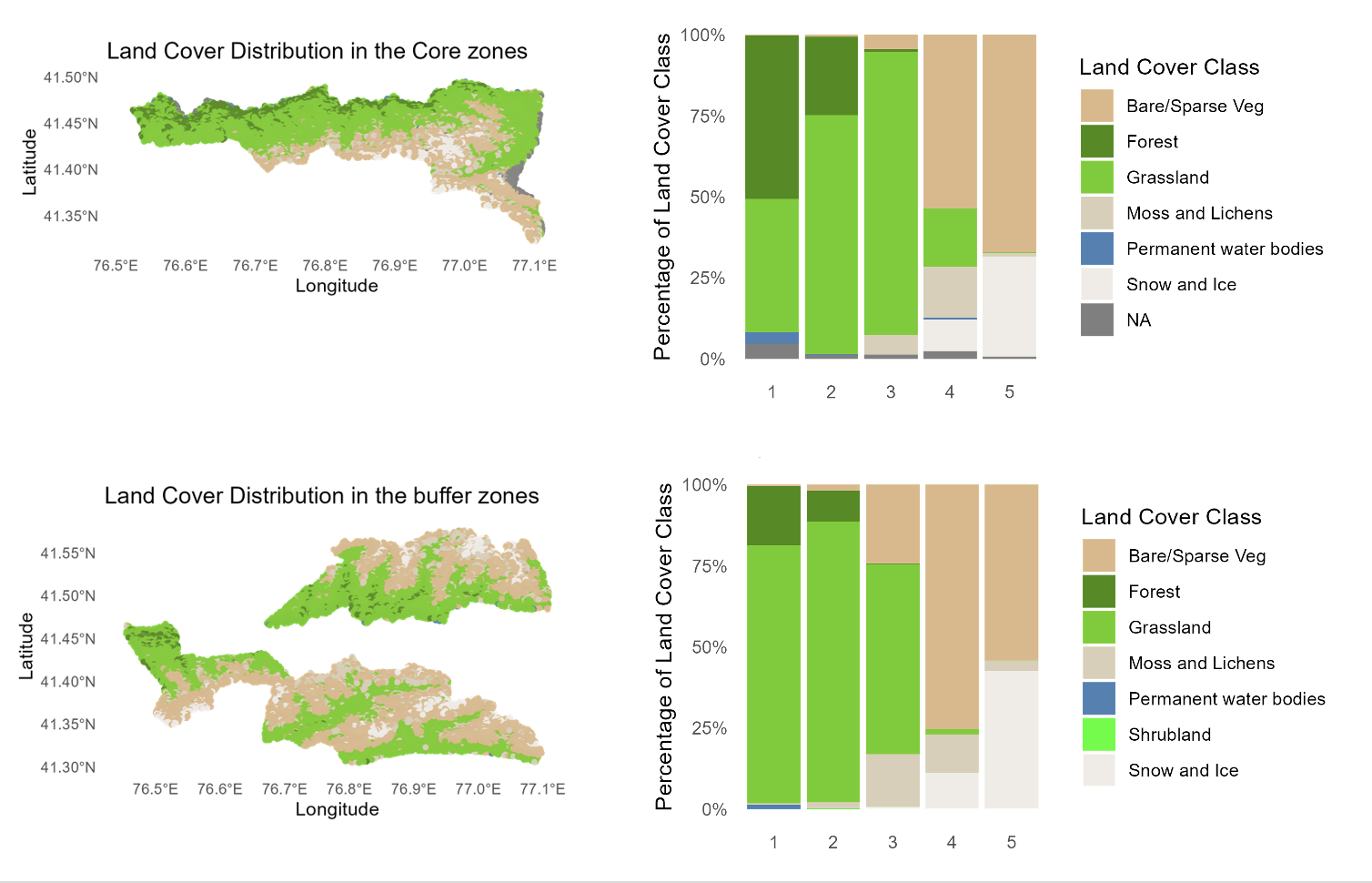

Figure A5. (Left column) Map of the land cover distribution across each study area extracted from the 2021 ESA Worldcover v200. (Right) Distribution of land cover proportion in each study area across elevation bands (See Table A1. in Appendix A for elevation band range).

Supervised Classification

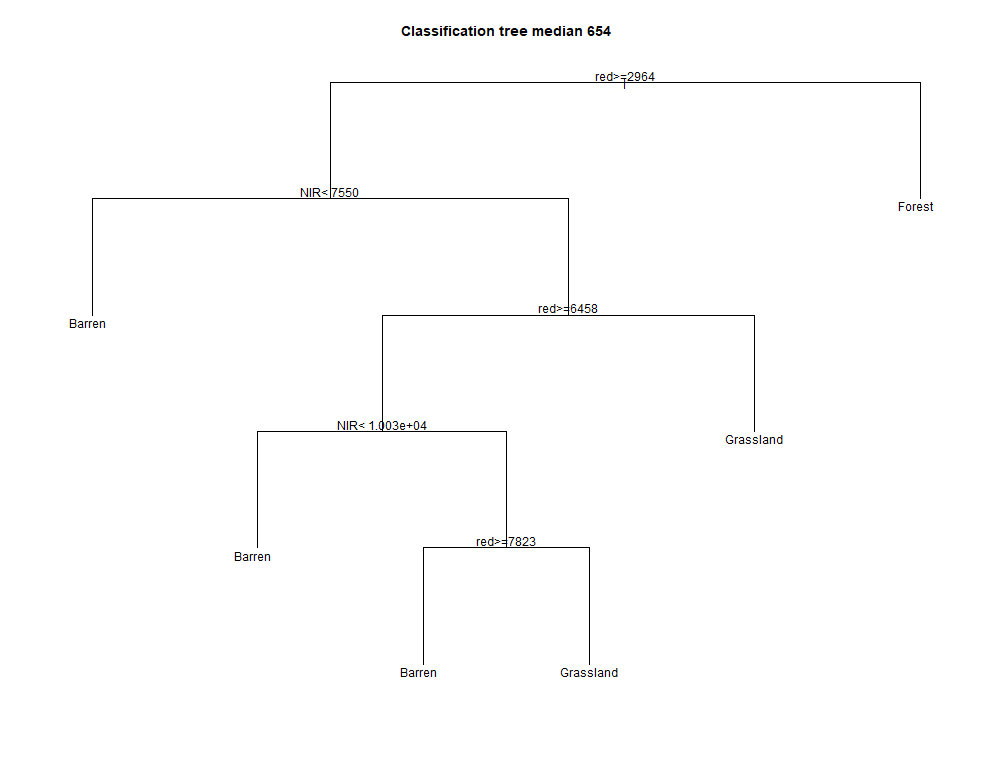

Figure A6.	Predicted land cover classes based on the surface reflectance for each pixel using the Classification and Regression Trees (CART) algorithm. The training dataset includes 1035 pixels for each land cover class to be classified. The spectral bands used to classify barren areas, forest and grasslands are the NIR, SWIR1 and red bands (median654 from GLAD Tools) and NIR, SWIR1 and SWIR2 (meidan567). We set a minimum of 200 observations (minsplit parameter) to constitute a class.

Table A3.	Confusion matrix representing the predicted and observed classification values of each given land cover class. The shaded diagonal represents the true positive classifications. The other values indicate the false positive classifications.

| 2021 | predicted | | |
| --- | --- | --- | --- |
| observed | Bare/Sparse Veg | Forest | Grassland |
| Barren | 978 | 24 | 33 |
| Forest | 18 | 896 | 121 |
| Grassland | 173 | 76 | 786 |

Table A4.	Classification accuracy of the Landsat multitemporal mosaic for 2017-2021 using 2021 ESA reference data. The user accuracy (UA) and producer accuracy (PA) are calculated per class and indicate the reliability of the classification and the capacity of the classifier to properly classify a pixel, respectively.

| 2021 | Accuracy | |
| --- | --- | --- |
| Land cover class | User Accuracy (UA) | Producer Accuracy (PA) |
| Barren | 0.84 | 0.95 |
| Forest | 0.91 | 0.87 |
| Grassland | 0.84 | 0.76 |

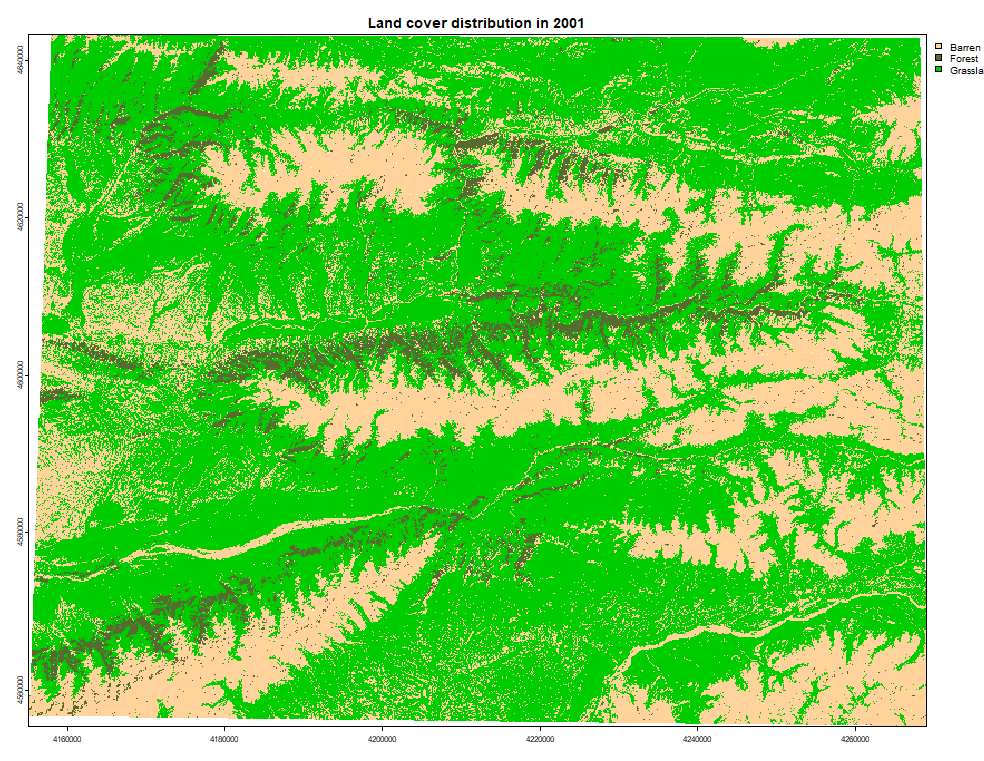

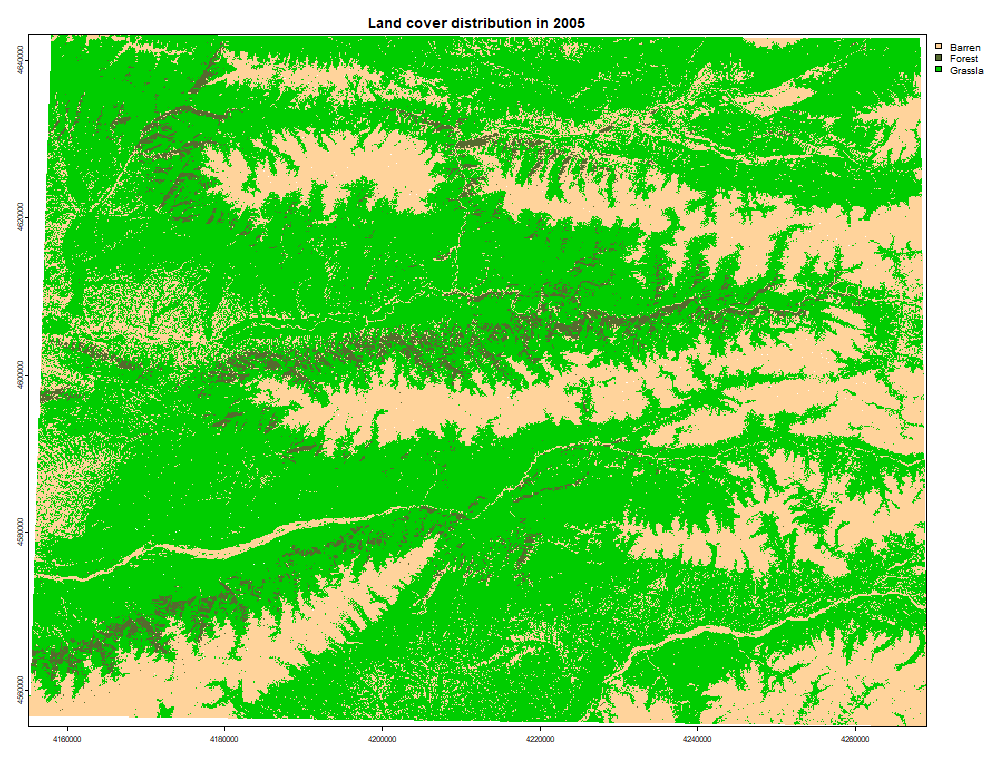

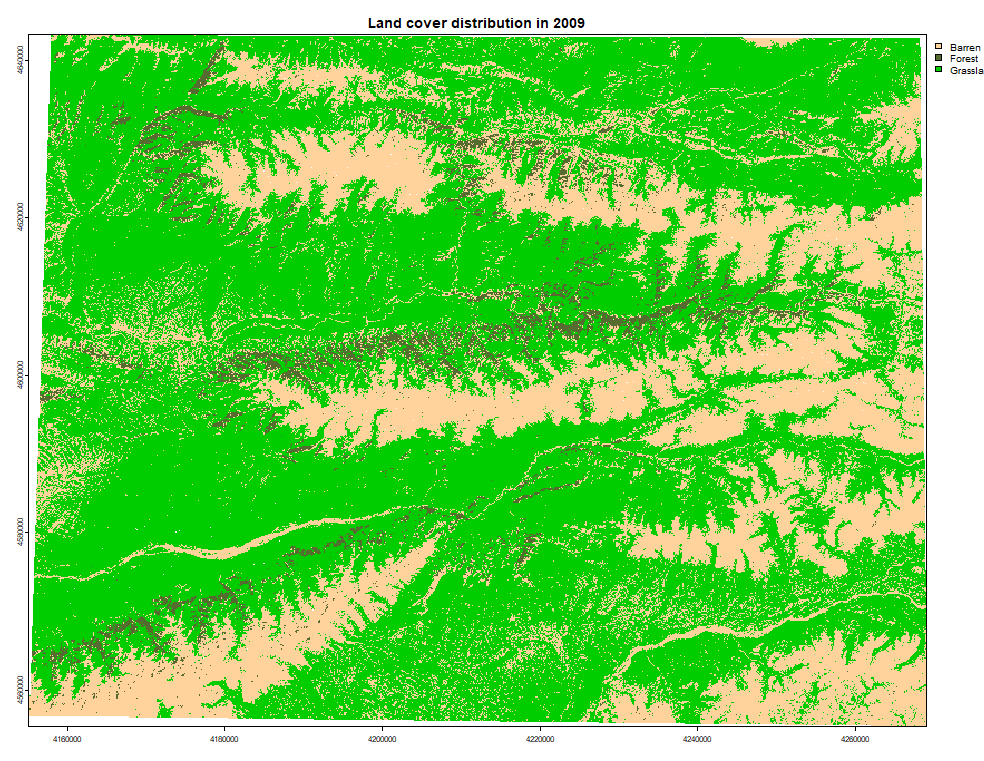

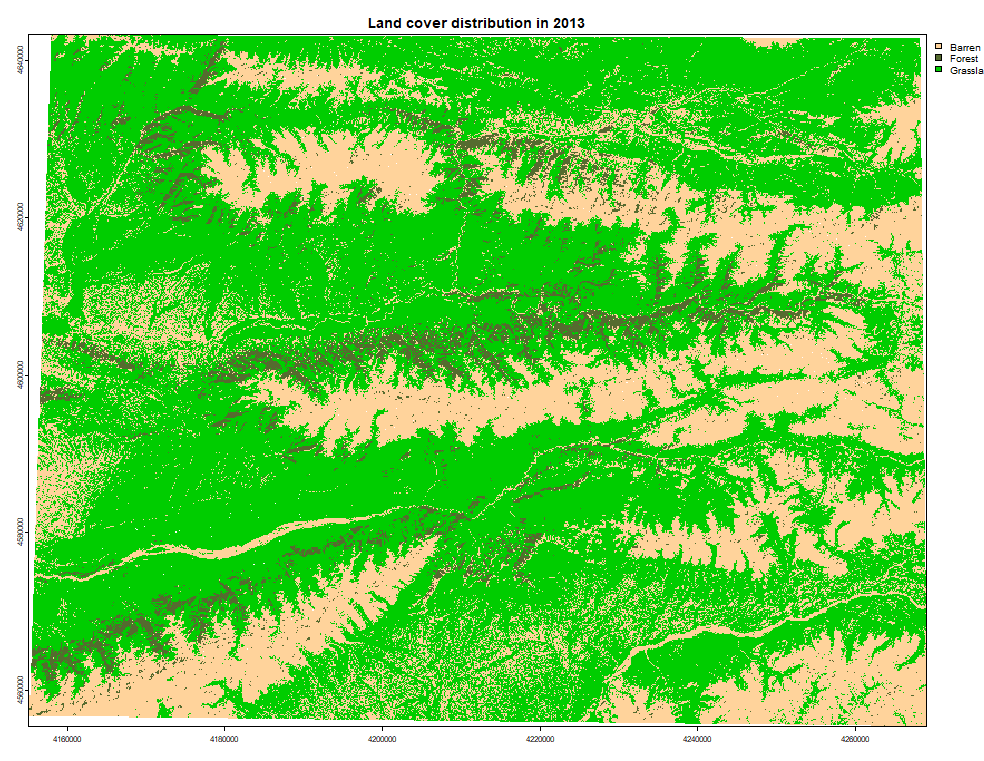

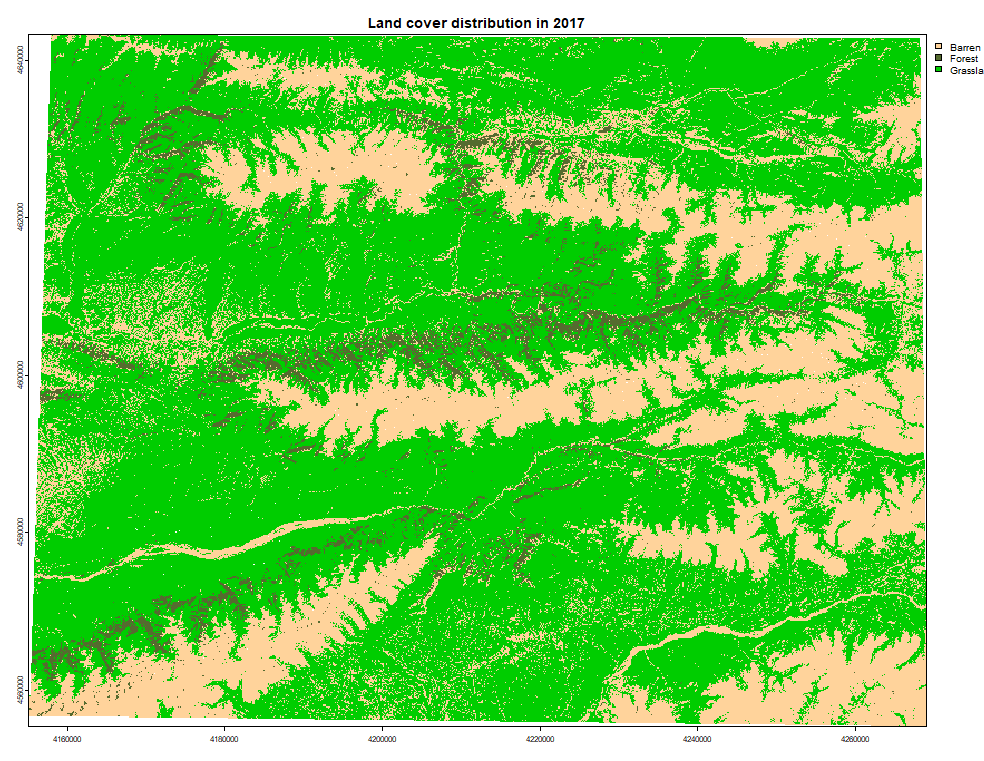

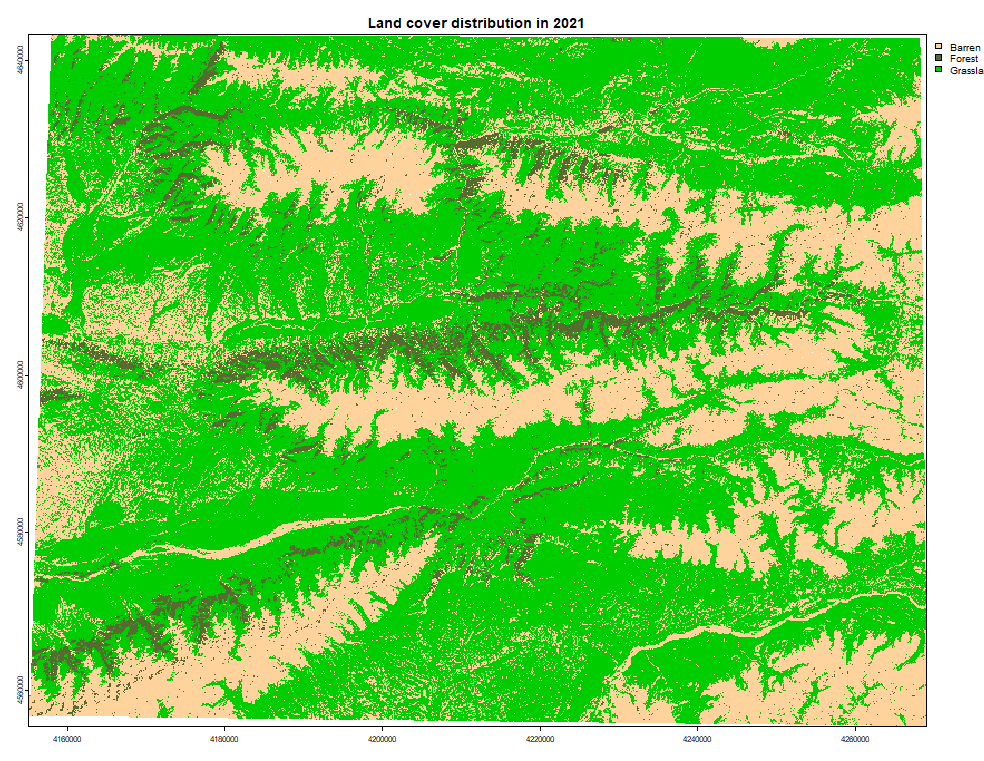

Figure A7.	Predicted grassland, barren, forest and snow/ice spatial distributions after the supervised classification of the Landsat multi-year metrics composite for each time period. (a) 1997-2001, (b) 2001-2005, (c) 2005-2009, (d) 2009-2013, (e) 2013-2017, (f) 2017-2021.

Boosted Regression Trees

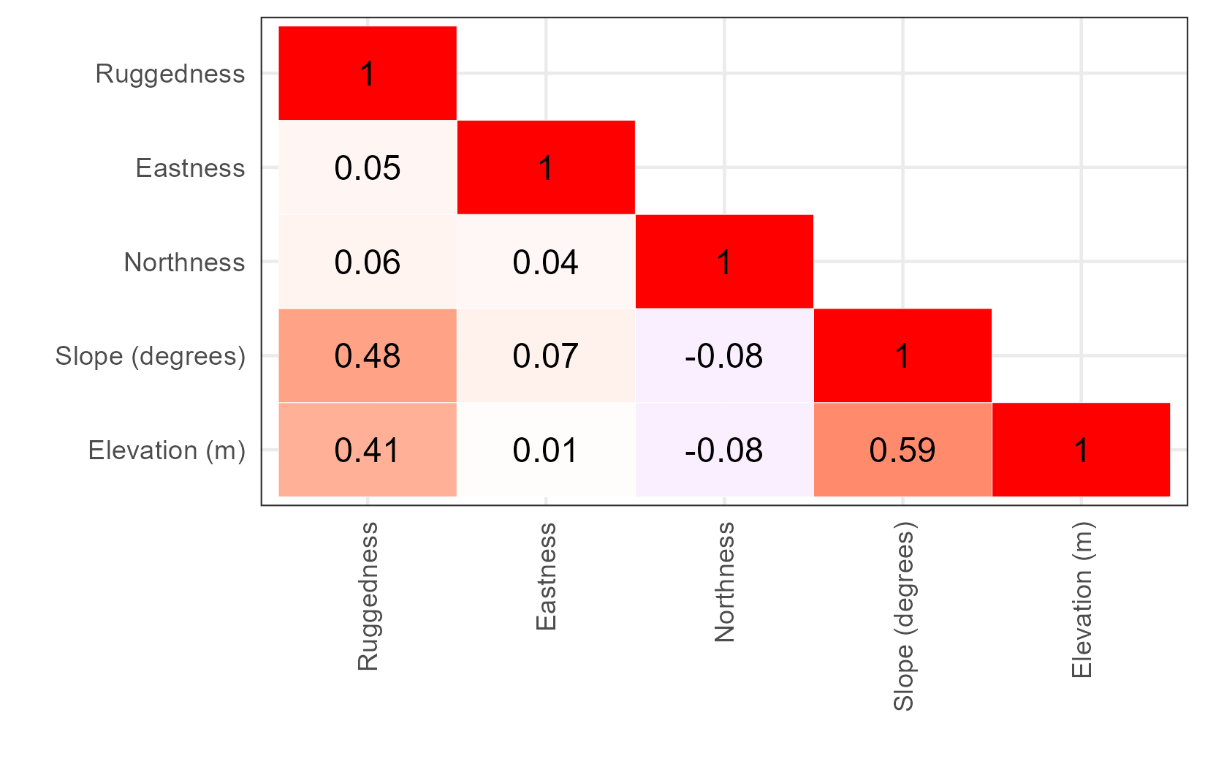

Figure A8.	Correlation matrix among explanatory variables from 3105 pixels.

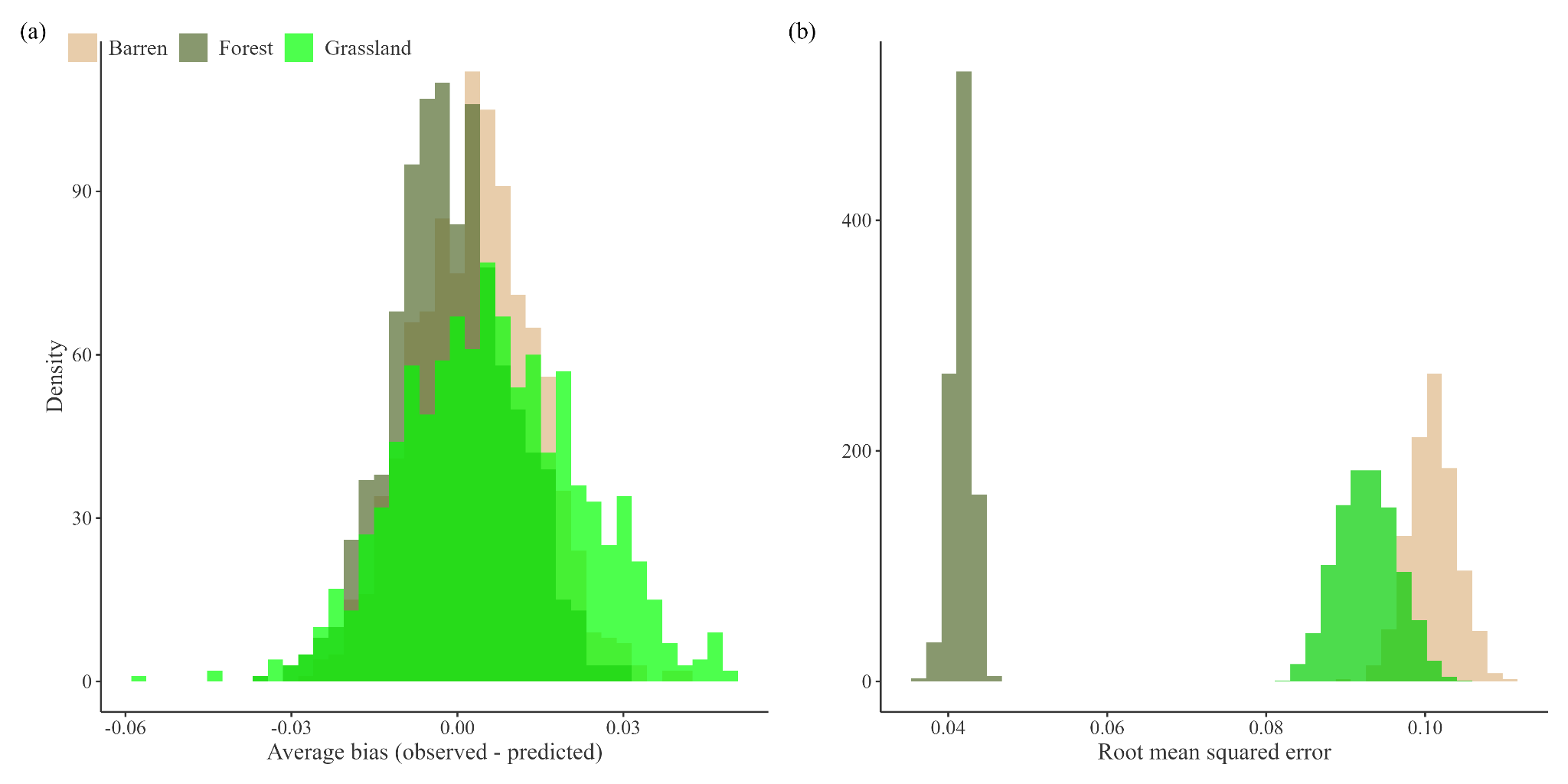

Figure A9.	(a) Average bias plot per land cover class. (b) Root mean squared error for each land cover class.

Table A5.	Selected hyperparameters for each land cover type. The minimization of the root mean square error selected in each model can be observed by comparing the selected lowest rmse (min_rmse) and the highest rmse (max_rmse) generated during the tuning step.

| Land cover | eta | gamma | max_depth | subsample | min_rmse | max_rmse |
| --- | --- | --- | --- | --- | --- | --- |
| Grassland | 0.5 | 0.7 | 10 | 0.3 | 0.1943 | 0.1970 |
| Barren | 0.2 | 0.5 | 15 | 0.3 | 0.0990 | 0.1024 |
| Forest | 0.5 | 0.5 | 10 | 0.9 | 0.0934 | 0.0957 |

Table A6.	Mean bias for each land cover class.

| Land cover | mean | sd | se | ci | lower | upper |
| --- | --- | --- | --- | --- | --- | --- |
| Barren | 0.00353 | 0.227 | 0.00119 | 0.00234 | 0.00119 | 0.00587 |
| Forest | -0.00096 | 0.223 | 0.00118 | 0.00231 | -0.00327 | 0.00135 |
| Grassland | 0.00636 | 0.337 | 0.00178 | 0.00348 | 0.00288 | 0.00984 |

Table A7. Mean root mean square error (rmse) for each land cover class.

| Land cover | mean | sd | se | ci | lower | upper |
| --- | --- | --- | --- | --- | --- | --- |
| Barren | 0.101 | 0.00300 | 0.00009 | 0.000186 | 0.101 | 0.101 |
| Forest | 0.0417 | 0.00134 | 0.00004 | 0.000083 | 0.0416 | 0.0417 |
| Grassland | 0.0927 | 0.000379 | 0.000120 | 0.000235 | 0.0924 | 0.0929 |

Table A8. Mean relative importance of each topographic variable on each land cover proportion.

| Land cover | Feature | mean | sd | lower | upper |
| --- | --- | --- | --- | --- | --- |
| Barren | Elevation | 0.561 | 0.0117 | 0.561 | 0.562 |
| Barren | Northness | 0.115 | 0.00749 | 0.114 | 0.115 |
| Barren | RRI | 0.113 | 0.00735 | 0.112 | 0.113 |
| Barren | Slope | 0.109 | 0.00748 | 0.109 | 0.109 |
| Barren | Eastness | 0.102 | 0.00708 | 0.102 | 0.102 |
| Forest | Elevation | 0.505 | 0.00844 | 0.504 | 0.505 |
| Forest | Northness | 0.296 | 0.00944 | 0.296 | 0.297 |
| Forest | Eastness | 0.0895 | 0.00654 | 0.0891 | 0.0899 |
| Forest | RRI | 0.0553 | 0.00447 | 0.0550 | 0.0556 |
| Forest | Slope | 0.0546 | 0.00655 | 0.0542 | 0.0550 |
| Grassland | Elevation | 0.265 | 0.0119 | 0.265 | 0.266 |
| Grassland | Northness | 0.206 | 0.0139 | 0.205 | 0.206 |
| Grassland | Slope | 0.194 | 0.0158 | 0.193 | 0.195 |
| Grassland | RRI | 0.173 | 0.0107 | 0.173 | 0.174 |
| Grassland | Eastness | 0.161 | 0.00982 | 0.161 | 0.162 |

Change detection

Table A9.	Transition matrix describing the raw number of pixels per land cover class across the three study areas between 2001 and 2021. For each land cover class, the total number of pixels that remained the same throughout the period is presented in the diagonal (shaded). The pixels that permanently transitioned from their initial 2001 classification (row) to another 2021 classification (column) are presented on either side of the diagonal. Each pixel is approximately 30 m x 30 m and our results only include pixels considered accurately classified (i.e. pixels obtained the same classification between the Landsat 2021 and the ESA; n = 54 847). The surface area characterized by each land cover is reported as percentages, as well as the proportion that changed.

| 2001 detection | 2021 detection | | |  | Total surface area (%) | Proportion changed  (%) |
| --- | --- | --- | --- | --- | --- | --- |
|  | Barren | Forest | Grassland | Total sites |  |  |
| Barren | 11997 | 1855 | 1095 | 17902 | 32.6% | 5.4% |
| Forest | 298 | 2142 | 120 | 2560 | 4.7% | 0.8% |
| Grassland | 3372 | 1606 | 29407 | 34385 | 62.7% | 9.1% |
| Surface area with the same land cover remaining (%) | | | | 84.8% |  |  |

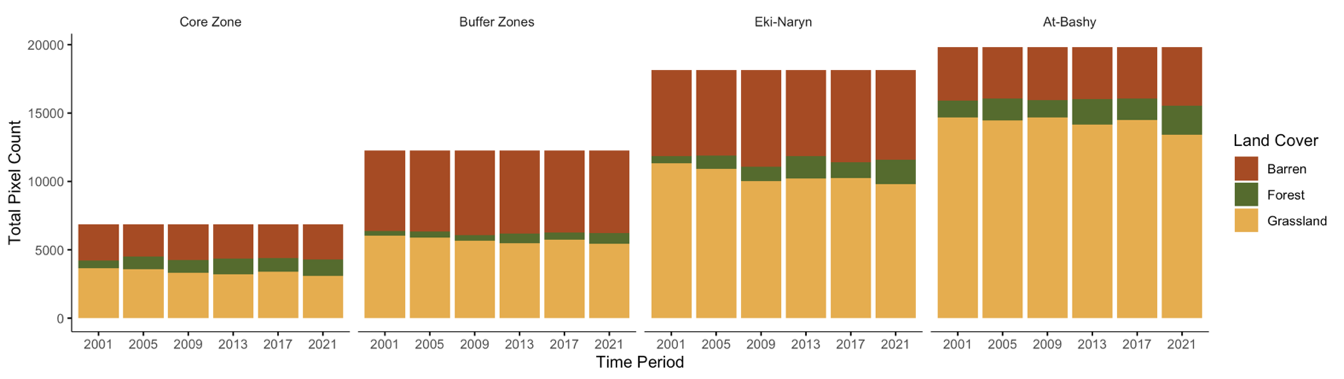

Figure A10.	Total numbers of pixels characterized as barren, forest or grassland in each given time period and study area. The study areas are organized from the least disturbed (core zone of Naryn State Reserve) to the most disturbed (At-Bashy). The land cover classification of each pixel was generated through the supervised classification of the 5-year Landsat composites.

Table A10.Percentage of the mean change between the main land cover transitions per (a) area, (b) period, (c) elevation. The direction of transitions between grasslands (G), barren (B) and forest (F) covers is indicated by the arrow.

|  | Mean percentage of change (%) | | |  |
| --- | --- | --- | --- | --- |
| (a) Area | G → B | B → G | G → F | B → F |
| Core | 4.20 | 3.01 | 1.72 | 0.02 |
| Buffer | 4.35 | 3.57 | 0.71 | 0.01 |
| Eki-Naryn | 5.08 | 4.13 | 0.48 | 0.02 |
| At-Bashy | 3.56 | 3.12 | 0.68 | 0.01 |
| (b) Time period | G → B | B → G | G → F | B → F |
| 2005 | 3.67 | 5.94 | 0.60 | 0.02 |
| 2009 | 6.52 | 4.26 | 0.71 | 0.01 |
| 2013 | 6.91 | 2.94 | 2.30 | 0.02 |
| 2017 | 4.09 | 3.49 | 0.31 | < 0.01 |
| 2021 | 4.59 | 4.11 | 1.48 | 0.02 |
| (c) Elevation band | G → B | B → G | G → F | B → F |
| 1 | 2.07 | 1.70 | 1.85 | 0.02 |
| 2 | 1.67 | 1.32 | 1.80 | 0.02 |
| 3 | 2.45 | 2.48 | 0.409 | 0.01 |
| 4 | 7.51 | 6.34 | 0.180 | 0.01 |
| 5 | 7.78 | 5.45 | 0.259 | 0.01 |

Raw data distribution

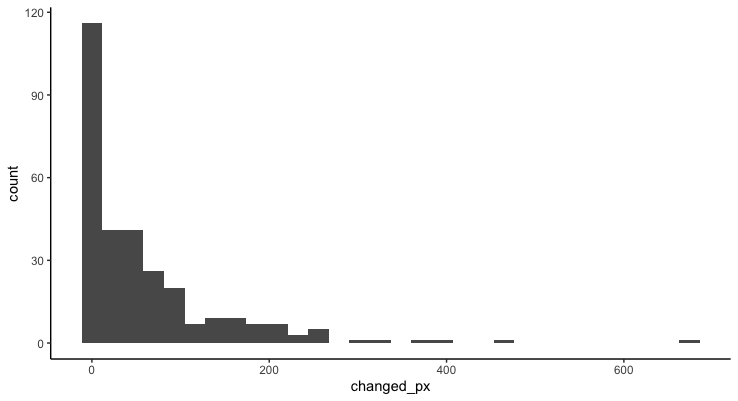

Figure A11.	Distribution of the count of pixels that changed between 2001 and 2021, regardless of the direction of change.

Model

Table A11.	Output from the generalized linear mixed model with negative binomial distribution showing the role of elevation, time, the degree of disturbance and the interaction between elevation and anthropogenic disturbance on the number of pixels that experienced browning or greening. To obtain proportions, we logged the total number of pixels, including those that did not experience change, and used the log total number of pixels as an offset. Significance codes represent 0 ‘***’, 0.001 ‘**’, 0.01 ‘*’, 0.05 ‘.’ and 1 ‘ ‘

| Predictors | Estimate | Std. Error | z value | Pr(>\|z\|) |  |
| --- | --- | --- | --- | --- | --- |
| (Intercept) | -3.973 | 0.25 | -15.874 | 0 | *** |
| elevation_band2 | 0.953 | 0.27 | 3.527 | 0 | *** |
| elevation_band3 | 1.087 | 0.267 | 4.064 | 0 | *** |
| elevation_band4 | 0.701 | 0.275 | 2.554 | 0.011 | * |
| elevation_band5 | -0.672 | 0.371 | -1.811 | 0.07 | . |
| transitionGreening | -0.621 | 0.275 | -2.258 | 0.024 | * |
| areaBuffer | 0.026 | 0.286 | 0.091 | 0.928 |  |
| areaEki-Naryn | 1.451 | 0.269 | 5.387 | 0 | *** |
| areaAt-Bashy | 0.584 | 0.269 | 2.172 | 0.03 | * |
| year2009 | 0.127 | 0.175 | 0.726 | 0.468 |  |
| year2013 | -0.461 | 0.174 | -2.644 | 0.008 | ** |
| year2017 | -0.939 | 0.176 | -5.32 | 0 | *** |
| year2021 | -0.043 | 0.177 | -0.243 | 0.808 |  |
| elevation_band2:transitionGreening | 0.15 | 0.231 | 0.648 | 0.517 |  |
| elevation_band3:transitionGreening | 0.245 | 0.229 | 1.069 | 0.285 |  |
| elevation_band4:transitionGreening | 0.11 | 0.235 | 0.467 | 0.64 |  |
| elevation_band5:transitionGreening | -0.117 | 0.338 | -0.345 | 0.73 |  |
| transitionGreening:areaBuffer | -0.023 | 0.222 | -0.103 | 0.918 |  |
| transitionGreening:areaEki-Naryn | -0.084 | 0.218 | -0.386 | 0.699 |  |
| transitionGreening:areaAt-Bashy | 0.018 | 0.219 | 0.083 | 0.934 |  |
| transitionGreening:year2009 | -0.286 | 0.249 | -1.149 | 0.251 |  |
| transitionGreening:year2013 | 0.314 | 0.246 | 1.275 | 0.202 |  |
| transitionGreening:year2017 | 1.02 | 0.246 | 4.148 | 0 | *** |
| transitionGreening:year2021 | -0.376 | 0.251 | -1.499 | 0.134 |  |
| elevation_band2:areaBuffer | -0.453 | 0.348 | -1.3 | 0.193 |  |
| elevation_band3:areaBuffer | -0.018 | 0.346 | -0.052 | 0.959 |  |
| elevation_band4:areaBuffer | -1.187 | 0.353 | -3.358 | 0.001 | *** |
| elevation_band5:areaBuffer | 0.219 | 0.476 | 0.459 | 0.646 |  |
| elevation_band2:areaEki-Naryn | -1.469 | 0.33 | -4.454 | 0 | *** |
| elevation_band3:areaEki-Naryn | -1.434 | 0.329 | -4.36 | 0 | *** |
| elevation_band4:areaEki-Naryn | -1.976 | 0.334 | -5.916 | 0 | *** |
| elevation_band5:areaEki-Naryn | -1.445 | 0.468 | -3.087 | 0.002 | ** |
| elevation_band2:areaAt-Bashy | -1.94 | 0.332 | -5.848 | 0 | *** |
| elevation_band3:areaAt-Bashy | -1.246 | 0.33 | -3.774 | 0 | *** |
| elevation_band4:areaAt-Bashy | -0.857 | 0.334 | -2.57 | 0.01 | * |
| elevation_band5:areaAt-Bashy | -1.408 | 0.48 | -2.932 | 0.003 | ** |

Pixel change across elevations and study areas

Table A12. Pairwise contrasts between the model-estimated marginal means in the proportion of pixels changed for different combinations of elevation and area. Estimated marginal means were calculated by marginalizing over levels of year and transition type (i.e., the contrasts represent the differences between elevation bands on average across years and transition types).

| contrast | area | estimate | SE | z.ratio | p.value |
| --- | --- | --- | --- | --- | --- |
| elevation_band1 - elevation_band2 | Core | -1.028 | 0.248 | -4.149 | < 0.001 |
| elevation_band1 - elevation_band3 | Core | -1.209 | 0.247 | -4.900 | < 0.001 |
| elevation_band1 - elevation_band4 | Core | -0.756 | 0.251 | -3.008 | 0.022 |
| elevation_band1 - elevation_band5 | Core | 0.730 | 0.346 | 2.108 | 0.216 |
| elevation_band2 - elevation_band3 | Core | -0.181 | 0.218 | -0.833 | 0.920 |
| elevation_band2 - elevation_band4 | Core | 0.272 | 0.223 | 1.219 | 0.740 |
| elevation_band2 - elevation_band5 | Core | 1.758 | 0.327 | 5.384 | < 0.001 |
| elevation_band3 - elevation_band4 | Core | 0.453 | 0.221 | 2.047 | 0.243 |
| elevation_band3 - elevation_band5 | Core | 1.939 | 0.325 | 5.960 | < 0.001 |
| elevation_band4 - elevation_band5 | Core | 1.486 | 0.328 | 4.533 | < 0.001 |
| elevation_band1 - elevation_band2 | Buffer | -0.575 | 0.245 | -2.345 | 0.131 |
| elevation_band1 - elevation_band3 | Buffer | -1.191 | 0.243 | -4.898 | < 0.001 |
| elevation_band1 - elevation_band4 | Buffer | 0.431 | 0.254 | 1.698 | 0.435 |
| elevation_band1 - elevation_band5 | Buffer | 0.511 | 0.333 | 1.533 | 0.541 |
| elevation_band2 - elevation_band3 | Buffer | -0.616 | 0.217 | -2.844 | 0.036 |
| elevation_band2 - elevation_band4 | Buffer | 1.006 | 0.228 | 4.411 | < 0.001 |
| elevation_band2 - elevation_band5 | Buffer | 1.086 | 0.315 | 3.444 | 0.005 |
| elevation_band3 - elevation_band4 | Buffer | 1.622 | 0.224 | 7.233 | < 0.001 |
| elevation_band3 - elevation_band5 | Buffer | 1.702 | 0.313 | 5.436 | < 0.001 |
| elevation_band4 - elevation_band5 | Buffer | 0.080 | 0.319 | 0.252 | 0.999 |
| elevation_band1 - elevation_band2 | Eki-Naryn | 0.441 | 0.217 | 2.031 | 0.251 |
| elevation_band1 - elevation_band3 | Eki-Naryn | 0.225 | 0.217 | 1.037 | 0.838 |
| elevation_band1 - elevation_band4 | Eki-Naryn | 1.220 | 0.220 | 5.533 | < 0.001 |
| elevation_band1 - elevation_band5 | Eki-Naryn | 2.175 | 0.320 | 6.798 | < 0.001 |
| elevation_band2 - elevation_band3 | Eki-Naryn | -0.216 | 0.214 | -1.011 | 0.850 |
| elevation_band2 - elevation_band4 | Eki-Naryn | 0.779 | 0.219 | 3.564 | 0.003 |
| elevation_band2 - elevation_band5 | Eki-Naryn | 1.734 | 0.319 | 5.445 | < 0.001 |
| elevation_band3 - elevation_band4 | Eki-Naryn | 0.995 | 0.219 | 4.553 | < 0.001 |
| elevation_band3 - elevation_band5 | Eki-Naryn | 1.950 | 0.318 | 6.129 | < 0.001 |
| elevation_band4 - elevation_band5 | Eki-Naryn | 0.955 | 0.320 | 2.988 | 0.024 |
| elevation_band1 - elevation_band2 | At-Bashy | 0.912 | 0.220 | 4.147 | < 0.001 |
| elevation_band1 - elevation_band3 | At-Bashy | 0.037 | 0.218 | 0.170 | 1.000 |
| elevation_band1 - elevation_band4 | At-Bashy | 0.101 | 0.219 | 0.461 | 0.991 |
| elevation_band1 - elevation_band5 | At-Bashy | 2.138 | 0.334 | 6.398 | < 0.001 |
| elevation_band2 - elevation_band3 | At-Bashy | -0.875 | 0.221 | -3.959 | 0.001 |
| elevation_band2 - elevation_band4 | At-Bashy | -0.811 | 0.224 | -3.623 | 0.003 |
| elevation_band2 - elevation_band5 | At-Bashy | 1.226 | 0.337 | 3.643 | 0.002 |
| elevation_band3 - elevation_band4 | At-Bashy | 0.064 | 0.220 | 0.290 | 0.998 |
| elevation_band3 - elevation_band5 | At-Bashy | 2.101 | 0.334 | 6.280 | < 0.001 |
| elevation_band4 - elevation_band5 | At-Bashy | 2.037 | 0.336 | 6.068 | < 0.001 |
